# Prioritizing 2nd and 3rd order interactions via support vector ranking using sensitivity indices on static Wnt measurements^†^

**DOI:** 10.1101/059469

**Authors:** shriprakash sinha

## Abstract

It is widely known that the sensitivity analysis plays a major role in computing the strength of the influence of involved factors in any phenomena under investigation. When applied to expression profiles of various intra/extracellular factors that form an integral part of a signaling pathway, the variance and density based analysis yields a range of sensitivity indices for individual as well as various combinations of factors. These combinations denote the higher order interactions among the involved factors that might be of interest in the working mechanism of the pathway. For example, *DACT*3 is known to be a epigenetic regulator of the Wnt pathway in colorectal cancer and subject to histone modifications. But many of the *n^th^* ≥ 2 order interactions of *DACT*3 that might be influential have not been explored/tested. In this work, after estimating the individual effects of factors for a higher order combination, the individual indices are considered as discriminative features. A combination, then is a multivariate feature set in higher order (≥ 2). With an excessively large number of factors involved in the pathway, it is difficult to search for important combinations in a wide search space over different orders. Exploiting the analogy of prioritizing webpages using ranking algorithms, for a particular order, a full set of combinations of interactions can then be prioritized based on these features using a powerful ranking algorithm via support vectors. The computational ranking sheds light on unexplored combinations that can further be investigated using hypothesis testing based on wet lab experiments. Here, the basic framework and results obtained on 2^*nd*^ and 3^*rd*^ order interactions for members of family of *DACT*, *SFRP*, *DKK* (to name a few) in both normal and tumor cases is presented using a static data set.

**Significance:** The search and wet lab testing of unknown biological hypotheses in the form of combinations of various intra/extracellular factors that are involved in a signaling pathway, costs a lot in terms of time, investment and energy. To reduce this cost of search in a vast combinatorial space, a pipeline has been developed that prioritises these list of combinations so that a biologist can narrow down their investigation. The pipeline uses kernel based sensitivity indices to capture the influence of the factors in a pathway and employs powerful support vector ranking algorithm. The generic workflow and future improvements are bound to cut down the cost for many wet lab experiments and reveal unknown/untested biological hypothesis.

## 1 Introduction

### 1.1 A short review

Sharma ^1^’s accidental discovery of the Wingless played a pioneering role in the emergence of a widely expanding research field of the Wnt signaling pathway. A majority of the work has focused on issues related to • the discovery of genetic and epigenetic factors affecting the pathway (Thorstensen *et al*. ^2^ & Baron and Kneissel ^3^), • implications of mutations in the pathway and its dominant role on cancer and other diseases (Clevers ^4^), • investigation into the pathway’s contribution towards embryo development (Sokol ^5^), homeostasis (Pinto *et al*. ^6^, Zhong *et al*. ^7^) and apoptosis (Pe´cina-Šlaus ^8^) and • safety and feasibility of drug design for the Wnt pathway (Kahn ^9^, Garber ^10^, Voronkov and Krauss ^11^, Blagodatski *et al*. ^12^ & Curtin and Lorenzi ^13^). Approximately forty years after the discovery, important strides have been made in the research work involving several wet lab experiments and cancer clinical trials (Kahn ^9^, Curtin and Lorenzi ^13^) which have been augmented by the recent developments in the various advanced computational modeling techniques of the pathway. More recent informative reviews have touched on various issues related to the different types of the Wnt signaling pathway and have stressed not only the activation of the Wnt signaling pathway via the Wnt proteins (Rao and Kühl ^14^) but also the on the secretion mechanism that plays a major role in the initiation of the Wnt activity as a prelude (Yu and Virshup ^15^).

The work in this paper investigates some of the current aspects of research regarding the pathway via sensitivity analysis while using static (Jiang *et al*. ^16^) data retrieved from colorectal cancer samples.

### 1.2 Canonical Wnt signaling pathway

Before delving into the problem statement, a brief introduction to the Wnt pathway is given here. From the recent work of Sinha ^17^, the canonical Wnt signaling pathway is a transduction mechanism that contributes to embryo development and controls homeostatic self renewal in several tissues (Clevers ^4^). Somatic mutations in the pathway are known to be associated with cancer in different parts of the human body. Prominent among them is the colorectal cancer case (Gregorieff and Clevers ^18^). In a succinct overview, the Wnt signaling pathway works when the Wnt ligand gets attached to the Frizzled(*FZD*)/*LRP* coreceptor complex. *FZD* may interact with the Dishevelled (*DVL*) causing phosphorylation. It is also thought that Wnts cause phosphorylation of the *LRP* via casein kinase 1 (*CK*1) and kinase *GSK*3. These developments further lead to attraction of Axin which causes inhibition of the formation of the degradation complex. The degradation complex constitutes of *AXIN*, the *β*-*catenin* transportation complex *APC*, *CK*1 and *GSK*3. When the pathway is active the dissolution of the degradation complex leads to stabilization in the concentration of *β*-*catenin* in the cytoplasm. As *β*-*catenin* enters into the nucleus it displaces the *GROUCHO* and binds with transcription cell factor *TCF* thus instigating transcription of Wnt target genes. *GROUCHO* acts as lock on *TCF* and prevents the transcription of target genes which may induce cancer. In cases when the Wnt ligands are not captured by the coreceptor at the cell membrane, *AXIN* helps in formation of the degradation complex. The degradation complex phosphorylates *β*-*catenin* which is then recognized by *FBOX*/*WD* repeat protein *β*-*TRCP*. *β*-*TRCP* is a component of ubiquitin ligase complex that helps in ubiquitination of *β*-*catenin* thus marking it for degradation via the proteasome. Cartoons depicting the phenomena of Wnt being inactive and active are shown in figures 1(A) and 1(B), respectively.

**Fig. 1.**
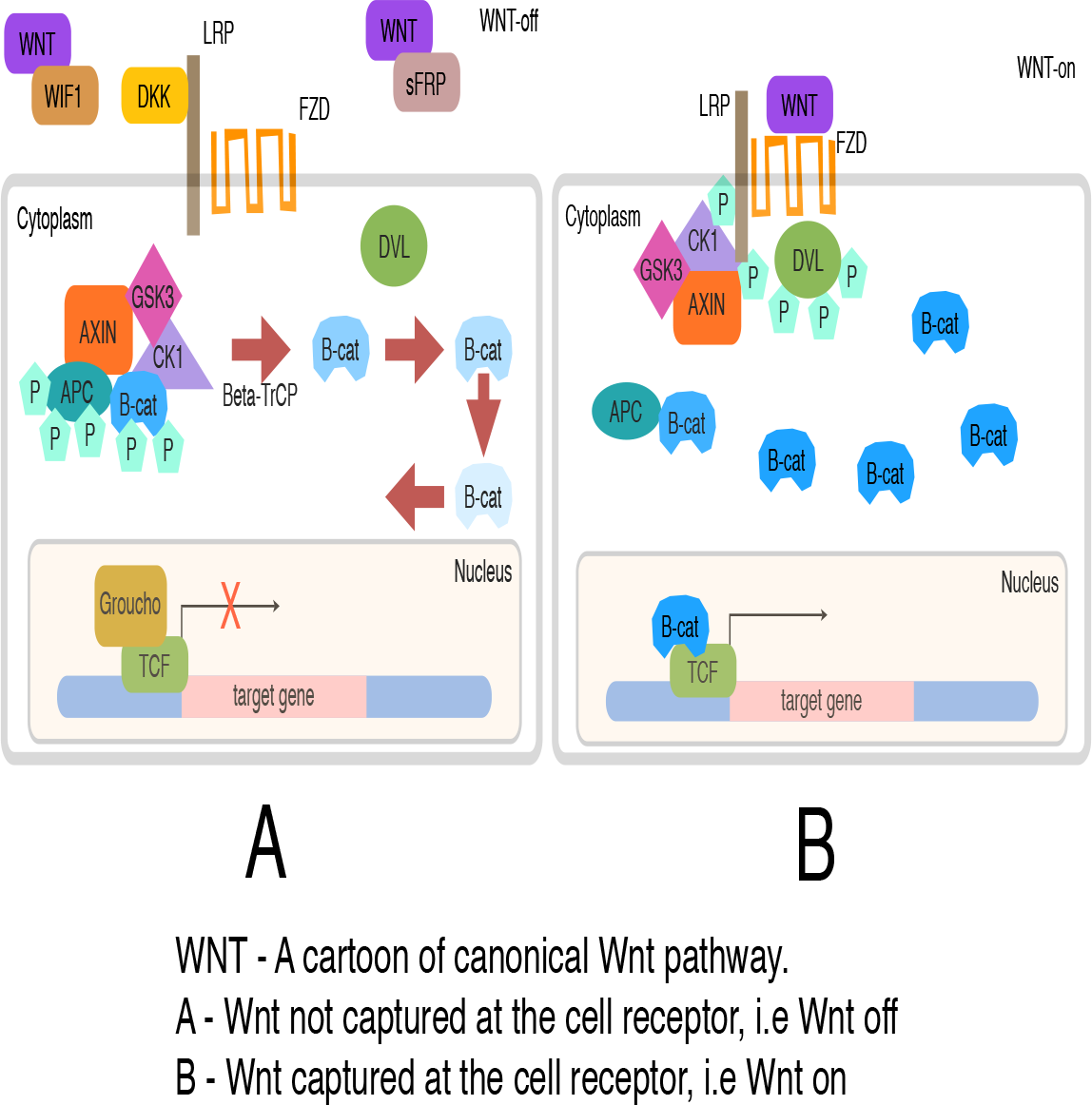
A cartoon of Wnt signaling pathway. Part (A) represents the destruction of *β*-*catenin* leading to the inactivation of the Wnt target gene. Part (B) represents activation of Wnt target gene.

**Fig. 2.**
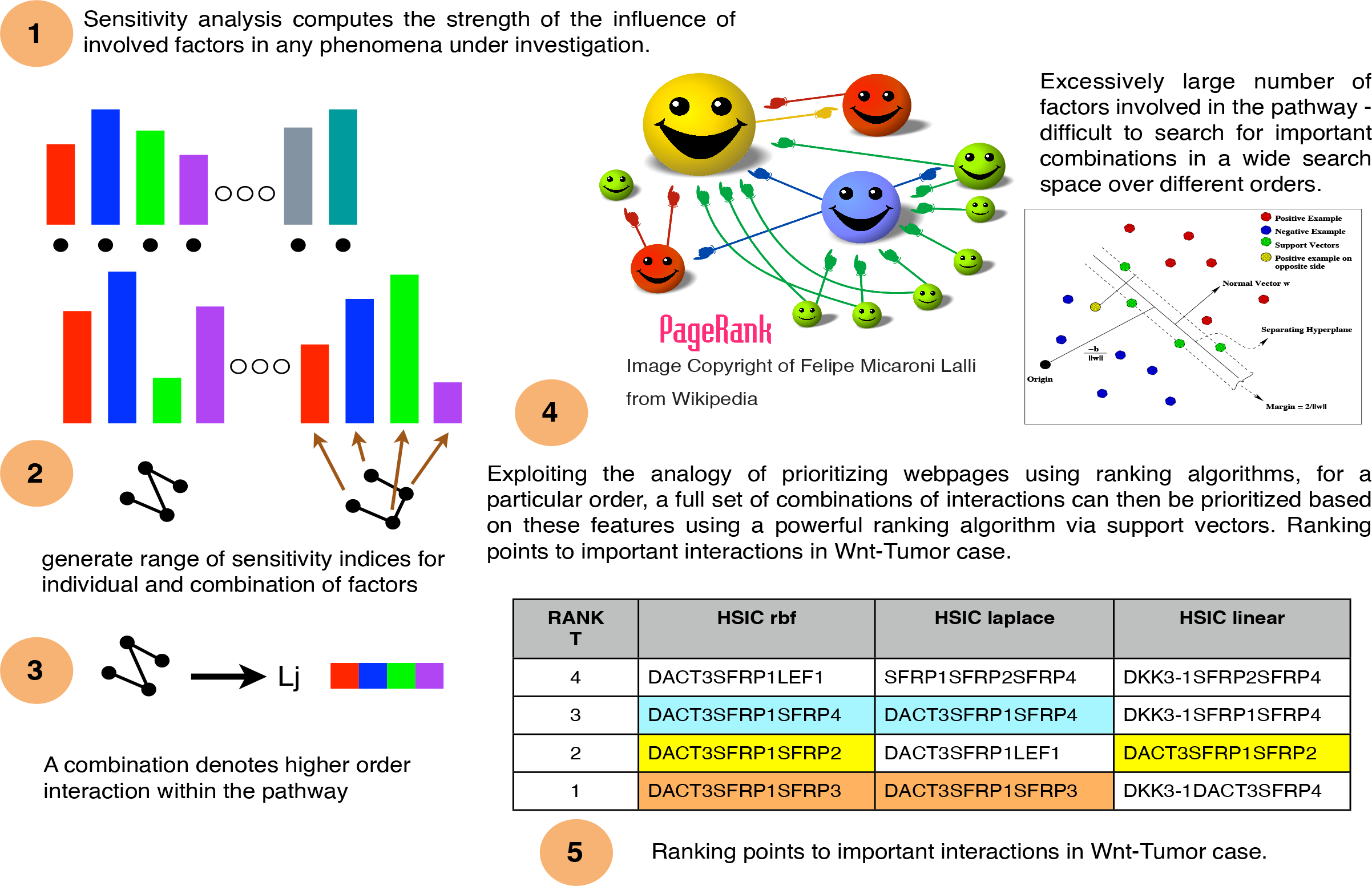
A graphical view of the general idea behind the current work. (1) Sensitivity indices capture the influence of involved factors in a pathway. (2) Generate sensitivity indices for individual and combinations of involved factors. Note that for a combination, the indices for the involved factors in the combination are generated separately. (3) Vectorize the indices per combination. (4) Rank these combinations based on the sensitivity indices using support vector ranking as web pages are ranked using a ranking algorithm. (5) Obtained is a prioritized list of interactions that could point to important interactions in the pathway in cancer cases.

## 2 Problem statement

It is widely known that the sensitivity analysis plays a major role in computing the strength of the influence of involved factors in any phenomena under investigation. When applied to expression profiles of various intra/extracellular factors that form an integral part of a signaling pathway, the variance and density based analysis yields a range of sensitivity indices for individual as well as various combinations of factors. These combinations denote the higher order interactions among the involved factors. Computation of higher order interactions is often time consuming but they give a chance to explore the various combinations that might be of interest in the working mechanism of the pathway. For example, in a range of fourth order combinations among the various factors of the Wnt pathway, it would be easy to assess the influence of the destruction complex formed by APC, AXIN, CSKI and GSK3 interaction. Unknown interactions can be further investigated by transforming biological hypothesis regarding these interactions in vitro, in vivo or in silico. But to mine these unknown interactions it is necessary to search a wide space of all combinations of input factors involved in the pathway.

In this work, after estimating the individual effects of factors for a higher order combination, the individual indices are considered as discriminative features. A combination, then is a multivariate feature set in higher order (>2). With an excessively large number of factors involved in the pathway, it is difficult to search for important combinations in a wide search space over different orders. Exploiting the analogy with the issues of prioritizing webpages using ranking algorithms, for a particular order, a full set of combinations of interactions can then be prioritized based on these features using a powerful ranking algorithm via support vectors (Joachims ^19^). The computational ranking sheds light on unexplored combinations that can further be investigated using hypothesis testing based on wet lab experiments. In this manuscript both local and global SA methods are used.

Similar higher order interactions can be computed and prioritized. This gives a chance to examine the biological hypothesis of interest regarding the positive/negative roles of unexplored combinations of the various factors involved in the Wnt pathway, in tumor and normal cases. Using recordings in time, it is possible to find how the prioritization of a specific combination of involved factors is changing in time. This further helps in revealing when an important combination like destruction complex formed by APC, AXIN, CSKI and GSK3 interaction could be influenced via an intervention. Thus a powerful computational mechanism is presented to explore the interactions involved in Wnt pathway. The framework can be used in any other pathway also.

## 3 Sensitivity analysis

Seminal work by Russian mathematician Sobol’ ^20^ lead to development as well as employment of SA methods to study various complex systems where it was tough to measure the contribution of various input parameters in the behaviour of the output. A recent unpublished review on the global SA methods by Iooss and Lemaître ^21^ categorically delineates these methods with the following functionality • screening for sorting influential measures (Morris ^22^ method, Group screening in Moon *et al*. ^23^ & Dean and Lewis ^24^, Iterated factorial design in Andres and Hajas ^25^, Sequential bifurcation in Bettonvil and Kleijnen ^26^ and Cotter ^27^ design), • quantitative indicies for measuring the importance of contributing input factors in linear models (Christensen ^28^, Saltelli *et al*. ^29^, Helton and Davis ^30^ and McKay *et al*. ^31^) and nonlinear models (Homma and Saltelli ^32^, Sobol ^33^, Saltelli ^34^, Saltelli *et al*. ^35^, Saltelli *et al*. ^36^, Cukier *et al*. ^37^, Saltelli *et al*. ^38^, & Tarantola *et al*. ^39^ Saltelli *et al*. ^40^, Janon *et al*. ^41^, Owen ^42^, Tissot and Prieur ^43^, Da Veiga and Gamboa ^44^, Archer *et al*. ^45^, Tarantola *et al*. ^46^, Saltelli and Annoni ^47^ and Jansen ^48^) and • exploring the model behaviour over a range on input values (Storlie and Helton ^49^ and Da Veiga *et al*. ^50^, Li *et al*. ^51^ and Hajikolaei and Wang ^52^). Iooss and Lemaître ^21^ also provide various criteria in a flowchart for adapting a method or a combination of the methods for sensitivity analysis. Figure 3 shows the general flow of the mathematical formulation for computing the indices in the variance based Sobol method. The general idea is as follows - A model could be represented as a mathematical function with a multidimensional input vector where each element of a vector is an input factor. This function needs to be defined in a unit dimensional cube. Based on ANOVA decomposition, the function can then be broken down into *ƒ*_0_ and summands of different dimensions, if *ƒ*_0_ is a constant and integral of summands with respect to their own variables is 0. This implies that orthogonality follows in between two functions of different dimensions, if at least one of the variables is not repeated. By applying these properties, it is possible to show that the function can be written into a unique expansion. Next, assuming that the function is square integrable variances can be computed. The ratio of variance of a group of input factors to the variance of the total set of input factors constitute the sensitivity index of a particular group. Detailed derivation is present in the Appendix.

**Fig. 3.**
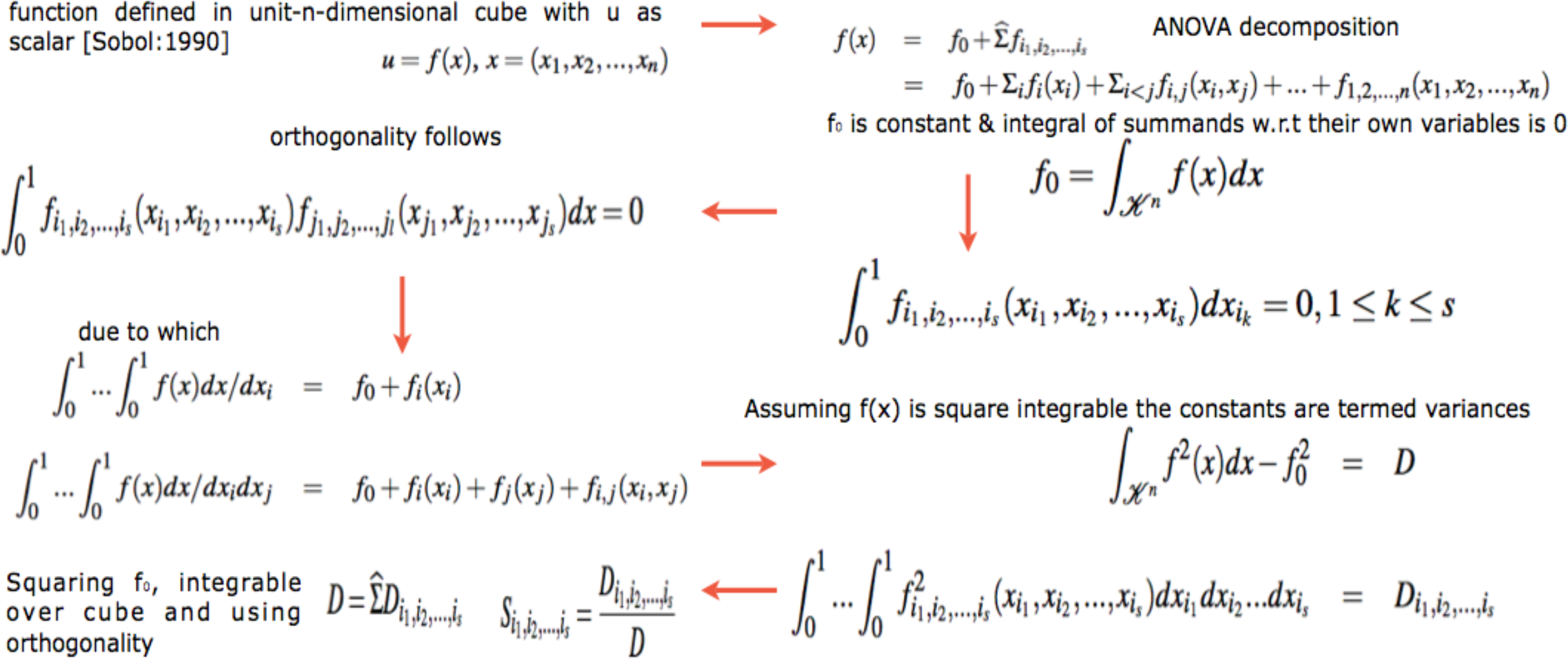
Computation of variance based sobol sensitivity indices. For detailed notations, see appendix.

Besides the above Sobol’ ^20^’s variance based indicies, more recent developments regarding new indicies based on density, derivative and goal-oriented can be found in Borgonovo ^53^, Sobol and Kucherenko ^54^ and Fort *et al*. ^55^, respectively. In a more recent development, Da Veiga ^56^ propose new class of indicies based on density ratio estimation (Borgonovo ^53^) that are special cases of dependence measures. This in turn helps in exploiting measures like distance correlation (Székely *et al*. ^57^) and Hilbert-Schmidt independence criterion (Gretton *et al*. ^58^) as new sensitivity indicies. The basic framework of these indicies is based on use of Csiszár *et al*. ^59^ f-divergence, concept of dissimilarity measure and kernel trick Aizerman *et al*. ^60^. Finally, Da Veiga ^56^ propose feature selection as an alternative to screening methods in sensitivity analysis. The main issue with variance based indicies (Sobol’ ^20^) is that even though they capture importance information regarding the contribution of the input factors, they • do not handle multivariate random variables easily and • are only invariant under linear transformations. In comparison to these variance methods, the newly proposed indicies based on density estimations (Borgonovo ^53^) and dependence measures are more robust. Figure 4 shows the general flow of the mathematical formulation for computing the indices in the density based HSIC method. The general idea is as follows - The sensitivity index is actually a distance correlation which incorporates the kernel based Hilbert-Schmidt Information Criterion between two input vectors in higher dimension. The criterion is nothing but the Hilbert-Schmidt norm of cross-covariance operator which generalizes the covariance matrix by representing higher order correlations between the input vectors through nonlinear kernels. For every operator and provided the sum converges, the Hilbert-Schmidt norm is the dot product of the orthonormal bases. For a finite dimensional input vectors, the Hilbert-Schmidt Information Criterion estimator is a trace of product of two kernel matrices (or the Gram matrices) with a centering matrix such that HSIC evalutes to a summation of different kernel values. Detailed derivation is present in the Appendix.

**Fig. 4.**
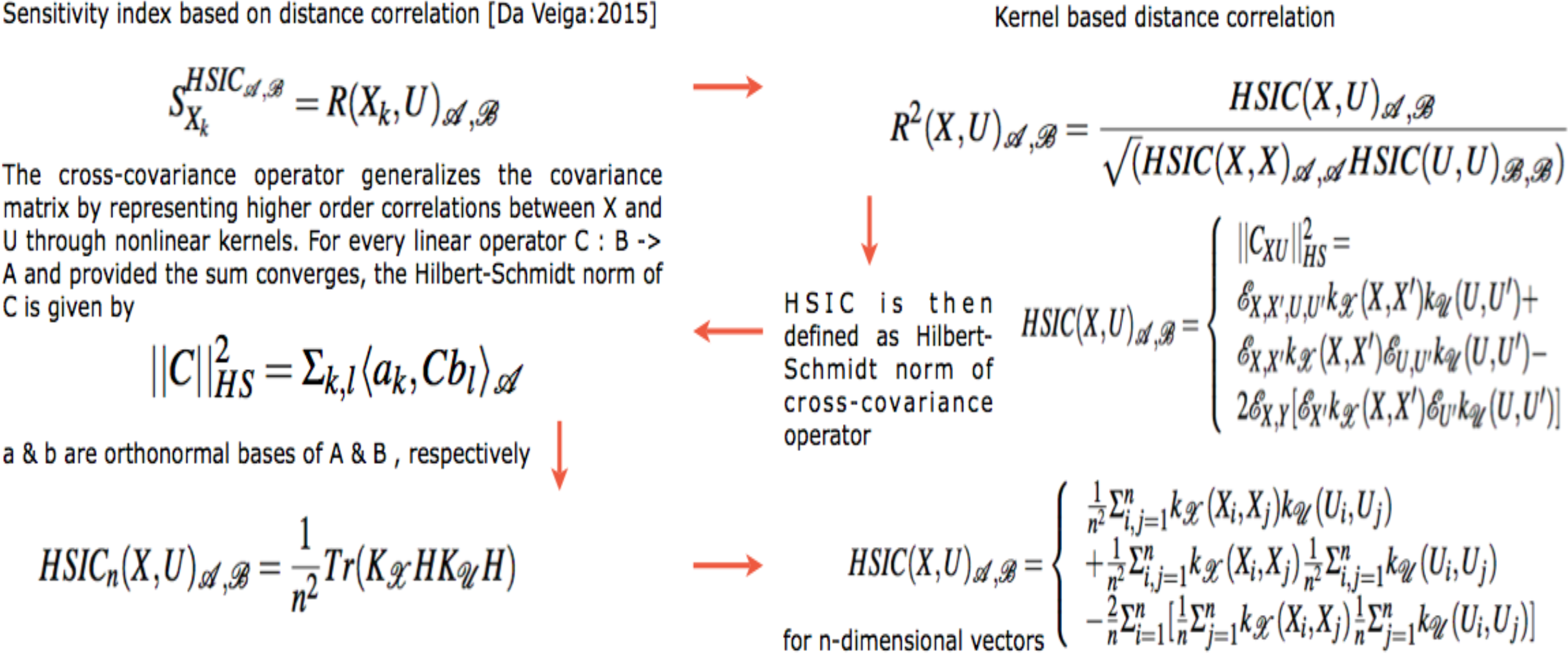
Computation of density based hsic sensitivity indices. For detailed notations, see appendix.

It is this strength of the kernel methods that HSIC is able to capture the deep nonlinearities in the biological data and provide reasonable information regarding the degree of influence of the involved factors within the pathway. Improvements in variance based methods also provide ways to cope with these nonlinearities but do not exploit the available strength of kernel methods. Results in the later sections provide experimental evidence for the same.

### 3.1 Relevance in systems biology

Recent efforts in systems biology to understand the importance of various factors apropos output behaviour has gained prominence. Sumner *et al*. ^61^ compares the use of Sobol’ ^20^ variance based indices versus Morris ^22^ screening method which uses a One-at-a-time (OAT) approach to analyse the sensitivity of *GSK*3 dynamics to uncertainty in an insulin signaling model. Similar efforts, but on different pathways can be found in Zheng and Rundell ^62^ and Marino *et al*. ^63^.

SA provides a way of analyzing various factors taking part in a biological phenomena and deals with the effects of these factors on the output of the biological system under consideration. Usually, the model equations are differential in nature with a set of inputs and the associated set of parameters that guide the output. SA helps in observing how the variance in these parameters and inputs leads to changes in the output behaviour. The goal of this manuscript is not to analyse differential equations and the parameters associated with it. Rather, the aim is to observe which input genotypic factors have greater contribution to observed phenotypic behaviour like a sample being normal or cancerous in both static and time series data. In this process, the effect of fold changes in time is also considered for analysis in the light of the recently observed psychophysical laws acting downstream of the Wnt pathway (Goentoro and Kirschner ^64^).

There are two approaches to sensitivity analysis. The first is the local sensitivity analysis in which if there is a required solution, then the sensitivity of a function apropos a set of variables is estimated via a partial derivative for a fixed point in the input space. In global sensitivity, the input solution is not specified. This implies that the model function lies inside a cube and the sensitivity indices are regarded as tools for studying the model instead of the solution. The general form of g-function (as the model or output variable) is used to test the sensitivity of each of the input factor (i.e expression profile of each of the genes). This is mainly due to its non-linearity, non-monotonicity as well as the capacity to produce analytical sensitivity indices. The g-function takes the form

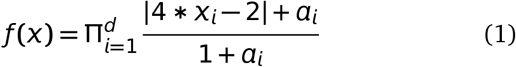

were, *d* is the total number of dimensions and *a_i_* **≥**0 are the indicators of importance of the input variable *X_i_*. Note that lower values of *a_i_* indicate higher importance of *X_i_*. In our formulation, we randomly assign values of *X_i_* [0, 1]. For the static (time series) data *d* **=** 18 (factors affecting the pathway). The value of *d* varies from 2 to 17, depending on the order of the combination one might be interested in. Thus the expression profiles of the various genetic factors in the pathway are considered as input factors and the global analysis conducted. Note that in the predefined dataset, the working of the signaling pathway is governed by a preselected set of genes that affect the pathway. For comparison purpose, the local sensitivity analysis method is also used to study how the individual factor is behaving with respect to the remaining factors while working of the pathway is observed in terms of expression profiles of the various factors.

Finally, in context of Goentoro and Kirschner ^64^’s work regarding the recent development of observation of Weber’s law working downstream of the pathway, it has been found that the law is governed by the ratio of the deviation in the input and the absolute input value. More importantly, it is these deviations in input that are of significance in studing such a phemomena. The current manuscript explores the sensitivity of deviation in the fold changes between measurements of fold changes at consecutive time points to explore in what duration of time, a particular factor is affecting the pathway in a major way. This has deeper implications in the fact that one is now able to observe when in time an intervention can be made or a gene be perturbed to study the behaviour of the pathway in tumorous cases. Thus sensitivity analysis of deviations in the mathematical formulation of the psychophysical law can lead to insights into the time period based influence of the involved factors in the pathway. Thus, both global and local anaylsis methods are employed to observe the entire behaviour of the pathway as well as the local behaviour of the input factors with respect to the other factors, respectively, via analysis of fold changes and deviations in fold changes, in time.

Given the range of estimators available for testing the sensitivity, it might be useful to list a few which are going to be employed in this research study. Also, a brief introduction into the fundamentals of the derivation of the three main indicies and the choice of sensitivity packages which are already available in literature, has been described in the Appendix.

## 4 Ranking Support Vector Machines

Learning to rank is a machine learning approach with the idea that the model is trained to learn how to rank. A good introduction to this work can be found in Li ^65^. Existing methods involve pointwise, pairwise and listwise approaches. In all these approaches, Support Vector Machines (SVM) can be employed to rank the required query. SVMs for pointwise approach build various hyperplanes to segregate the data and rank them. Pairwise approach uses ordered pair of objects to classify the objects and then utilize the classifyer to rank the objects. In this approach, the group structure of the ranking is not taken into account. Finally, the listwise ranking approach uses ranking list as instances for learning and prediction. In this case the ranking is straightforward and the group structure of ranking is maintained. Various different designs of SVMs have been developed and the research in this field is still in preliminary stages. In context of the gene expression data set employed in this manuscript, the objects are the genes with their RECORDED EXPRESSION VALUES FOR NORMAL AND TUMOR CASES. Both cases are treated separately.

Note that rankings algorithms have been developed to be employed in the genomic datasets but to the author’s awareness, these algorithms do not rank the range of combinations in a wide combinatorial search space in time. Also, they do not take into account the ranking of unexplored biological hypothesis which are assigned to a particular sensitivity value or vector that can be used for prioritization. For example, Kolde *et al*. ^66^ presents a ranking algorithm that betters existing ranking model based on the assignment of *P*-value. As stated by Kolde *et al*. ^66^ it detects genes that are ranked consistently better than expected under null hypothesis of uncorrelated inputs and assigns a significance score for each gene. The underlying probabilistic model makes the algorithm parameter free and robust to outliers, noise and errors. Significance scores also provide a rigorous way to keep only the statistically relevant genes in the final list. The proposed work here develops on sensitivity analysis and computes the influences of the factors for a system under investigation. These sensitivity indices give a much realistic view of the biological influence than the proposed *P*-value assignment and the probabilistic model. The manuscript at the current stage does not compare the algorithms as it is a pipeline to investigate and conduct a systems wide study. Instead of using SVM-Ranking it is possible to use other algorithms also, but the author has restricted to the development of the pipeline per se. Finally, the current work tests the effectiveness of the variance based (SOBOL) sensitivity indices apropos the density and kernel based (HSIC) sensitivity indices. Finally, Blondel *et al*. ^67^ provides a range of comparison for 10 different regression methods and a score to measure the models. Compared to the frame provided in Blondel *et al*. ^67^, the current pipeline takes into account biological information an converts into sensitivity scores and uses them as discriminative features to provide rankings. Thus the proposed method is algorithm independent.

## 5 Description of the dataset & design of experiments

A simple static dataset containing expression values measured for a few genes known to have important role in human colorectal cancer cases has been taken from Jiang *et al*. ^16^. Most of the expression values recorded are for genes that play a role in Wnt signaling pathway at an extracellular level and are known to have inhibitory affect on the Wnt pathway due to epigenetic factors. For each of the 24 normal mucosa and 24 human colorectal tumor cases, gene expression values were recorded for 14 genes belonging to the family of *SFRP*, *DKK*, *WIF*1 and *DACT*. Also, expression values of established Wnt pathway target genes like *LEF*1, *MYC*, *CD*44 and *CCND*1 were recorded per sample.

Note that green (red) represents activation (repression) in the heat maps of data in Jiang *et al*. ^16^. For the static data, only the scaled results are reported. This is mainly due to the fact that the measurements vary in a wide range and due to this there is often an error in the computed estimated of these indices. Bootstrapping without replicates on a smaller sample number is employed to generate estimates of indices which are then averaged. This takes into account the variance in the data and generates confithe dataset is prepared in the required format from both normal dence bands for the indices.

GENERAL ISSUES - • Here the input factors are the gene expression values for both normal and tumor cases in static data. For the case of time series data, the input factors are the fold change (deviations in fold change) expression values of genes at different time points (periods). Also, for the time series data, in the first experiment the analysis of a pair of the fold changes recorded at to different consecutive time points i.e *t_i_* &*t_i_*_**+**1_ is done. In the second experiment, the analysis of a pair of deviations in fold changes recorded at *t_i_* & *t_i_*_**+**1_ and *t_i_*_**+**1_ & *t_i_*_**+**2_. In this work, in both the static and the time series datasets, the analysis is done to study the entire model/pathway rather than find a particular solution to the model/pathway. Thus global sensitivity analysis is employed. But the local sensitivity methods are used to observe and compare the affect of individual factors via 1^*st*^ order analysis w.r.t total order analysis (i.e global analysis). In such an experiment, the output is the sensitivity indices of the individual factors participating in the model. This is different from the general trend of observing the sensitivity of parameter values that affect the pathway based on differential equations that model a reaction. Thus the model/pathway is studied as a whole by observing the sensitivities of the individual factors.

Note that the 24 normal and tumor cases are all different from each other. The 18 genes that are used to study in ^16^ are the input factors and it is unlikely that there will be correlations between different patients. The phenotypic behaviour might be similar at a grander scale. Also, since the sampling number is very small for a network of this scale, large standard deviations can be observed in many results, especially when the Sobol method is used. But this is not the issue with the sampling number. By that analysis, large deviations are not observed in kernel based density methods. The deviations are more because of the fact that the nonlinearities are not captured in an efficient way in the variance based Sobol methods. Due to this, the resulting indicies have high variance in numerical value. For the same number of samplings, the kernel based methods don’t show high variance.

The procedure begins with the listing of all 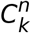 combinations for *k* number of genes from a total of *n* genes. *k* is ≥ 2 and ≤ (*n* − 1). Each of the combination of order *k* represent a unique set of interaction between the involved genetic factors. While studying the interaction among the various genetic factors using static data, tumor samples are considered separated from normal samples. For the experiments conducted here on a toy example, 20 bootstrap replicates were generated for each of normal and tumor samples without replacement. For each bootstrap replicate, the normal and turmor samples are divided into two different sets of equal size. Next the datasets are combined in a specifed format which go as input as per the requirement of a particular sensitivity analysis method. Thus for each *p^th^* combination in 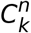 combinations, the dataset is prepared in the required format from both normal and tumor samples (See .R code in mainscript-1-1.R in google drive and step 3 in figure 5). After the data has been transformed, vectorized programming is employed for density based sensitivity analysis and looping is employed for variance based sensitivity analysis to compute the required sensitivity indices for each of the *p* combinations. Once the sensitivity indices are generated for each of the *p^th^* combination, for every bootstrap replicate in normal and tumor cases, confidence intervals are estimated for each sensitivity index. This procedure is done for different kinds of sensitivity analysis methods.

**Fig. 5.**
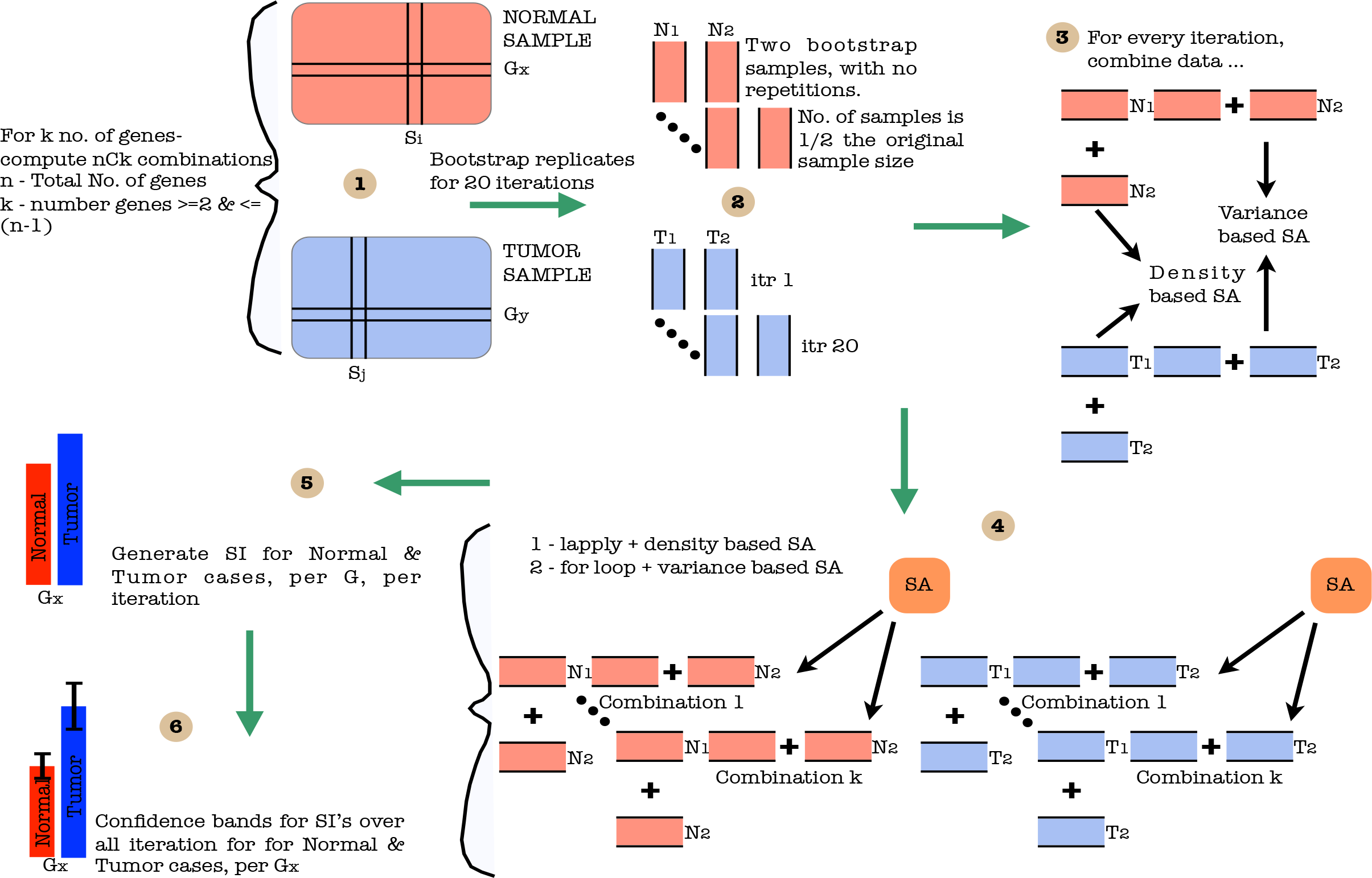
A cartoon of experimental setup. IMPORTANT NOTE - In this figure, *G_X_*, *G_y_* and *G* represent a combination. Step - (1) Segregation of data into normal and tumor cases. (2) Further data division per case and bootstrap sampling with no repetitions for different iterations. (3) Assembling bootstrapped data and application of SA methods. (4) Generation of SI’s for normal and tumor case per gene per iteration. (5) Generation of averaged SI and confidence bands per case per gene.

After the above sensitivity indices have been stored for each of the *p^th^* combination, each of the sensitivity analysis method for normal and tumor cases per bootstrap replicates, the next step in the design of experiment is conducted. Here, for a particular *k^th^* order of combination, a choice is made regarding the number of sample size (say *p*), where 2 ≤ *p ≤ noObs–* 1 (noObs is the number of observations i.e 20 replicates). Then for all sample sets of order *p* in 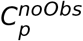, generate training index set of order *p* and test index set of order *noObs* − *p*. For each of the sample set, considering the sensitivity index for each individual factor of a gene combination in the previous step as described in the foregoing paragraph, a training and a test set is generated. Thus an observation in a training and a test set represents a gene combination with sensitivity indices of involved genetic factors as feature values. For a particular gene combination there are *p* training samples *noObs p* test samples. In total there are 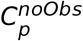 training sets and corresponding test sets. Next, 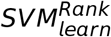 (Joachims ^19^) is used to generate a model on default value *C* value of 20. In the current experiment on toy model *C* value has not been tunned. The training set helps in the generation of the model as the different gene combinations are numbered in order which are used as rank indices. The model is then used to generate score on the observations in the testing set using the 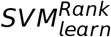(Joachims ^19^). Next the scores are averaged across all 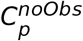 test samples. The experiment is conducted for normal and tumor samples separately. This procedure is executed for each and every sensitivity analysis method. Finally, for each sensitivity analysis method, for all *k^th^* order combinations, the mean across the averaged *p* scores is computed. This is followed by sorting of these scores along with the rank indices already assigned to the gene combinations. The end result is a sorted order of the gene combinations based on the ranking score learned by the *SVM^Rank^* algorithm. These steps are depicted in figure 6.

**Fig. 6.**
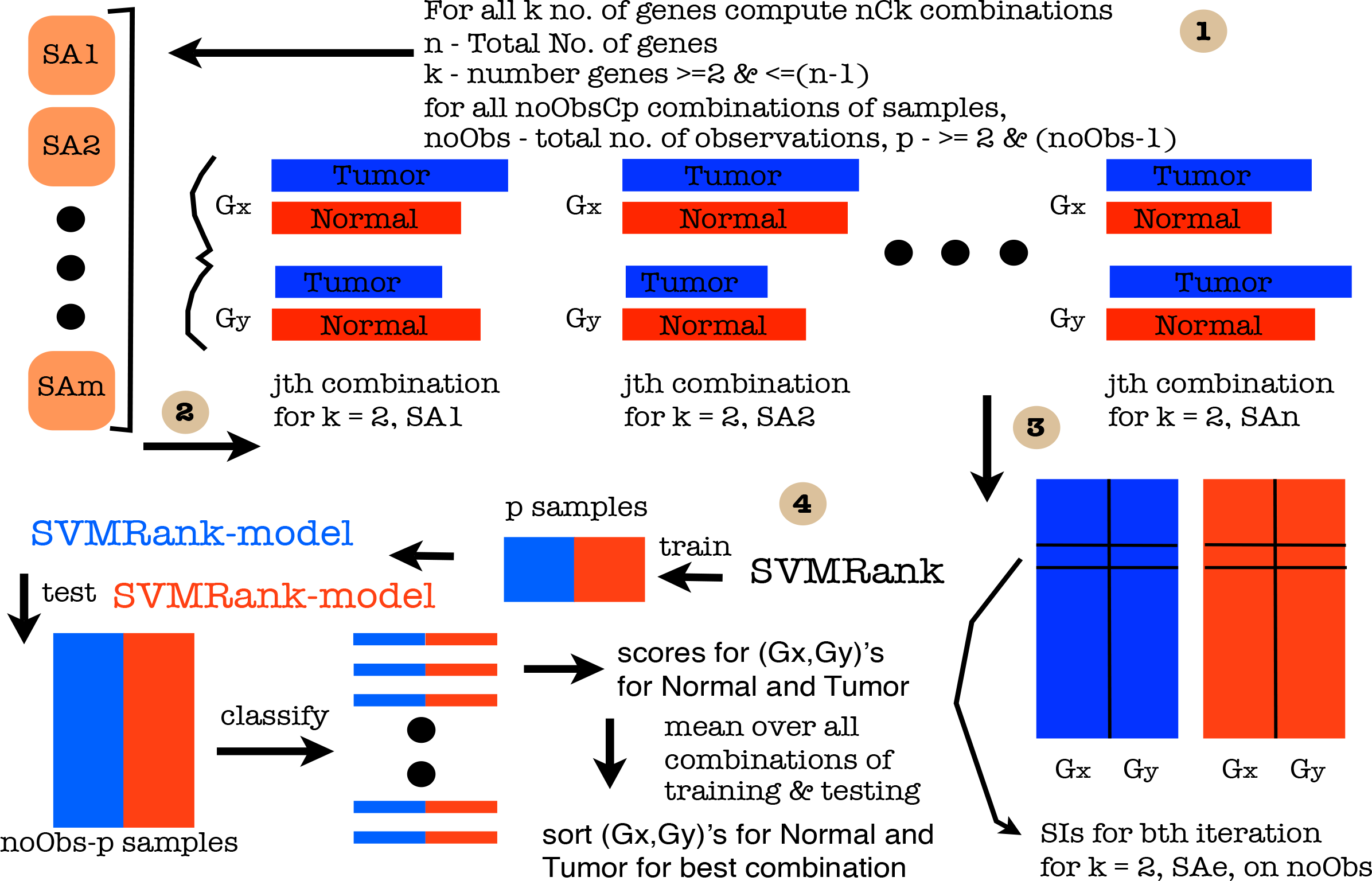
A cartoon of experimental setup. IMPORTANT NOTE - In this figure, *G_X_*, *G_y_* and *G* represent separate genes. Step - (1) Assembling *p* training indices and *noObs* − *p* testing indices for every *p^th^* order of samples in 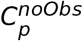 Thus there are a total of 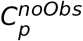training and corresponding test sets. (2) For every SA, combine (say for *k* **=** 2, i.e interaction level 2) SI’s of genetic factors for normal and tumor separately, for each observation in training and test data. (3) For *noObs* **=** 20 different replicates, per *SA_e_* and a particular combination of *< G_X_, G_y_ >* in normal and tumor, a matrix of observations is consturcted. (4) Using indices in (1)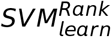 is employed on *p* training data to generate a model. This model is used to generate a ranking score on the test data via 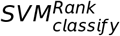. These score are averaged over 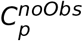 test data sets. Further, mean of scores over *noObs* − *p* test replicates per *< G_X_, G_y_ >* are computed and finally the combinations are ranked based on sorting for each of normal and training set.

Note that the following is the order in which the files should be executed in R, in order, for obtaining the desired results (Note that the code will not be explained here) - ● use source("mainScript-1-1.R") with arguments for Static data ● use source("Combine-Static-files.R"), if computing indices separately via previous file, ● source("Store-Results-S.R"), source("SVMRank-Results-S.R"), again this needs to be done separately for different kinds of SA methods and ● source("Sort-n-Plot-S.R") to sort the interactions.

## 6 Results and Discussion

Initial results on prioritization of 2^nd^ order interactions learnt from support vector ranking using these sensitivity indices were plotted for visualization. For a training sample size of 4 and test sample size of 16, from a total of 20 bootstraps for each of the interactions, in normal and tumor cases, and the sensitivity index using *Fdiv – Chi*2, the computed sensitivity indices were sorted in increasing order of value and are shown in figure 7 and 8, respectively. The *y – aXis* represents the sensitivity indices and the *X – aXis* represents the values of the different factors involved under investigation (here 2^*nd*^ order combinations). Since the indices have been sorted based on the ranking score, the combinations are now ordered according to the strength of their significance. The combinations with low significance or sensitivity index is placed towards the left most end and vice versa. Visualizations for other indices were also generated and are relegated to the appendix. Since various indices have been tested, it would be important to see how good the rankings tally across the different sensitivity methods. To tally the rankings visually, the top and the bottom 10 combinations were tabulated in tables 1, 2, 5, 3 and 4. The top and the bottom 10 rankings in figures 7 and 8 can be found in the second columns of tables 1 and 2.

**Fig. 7.**
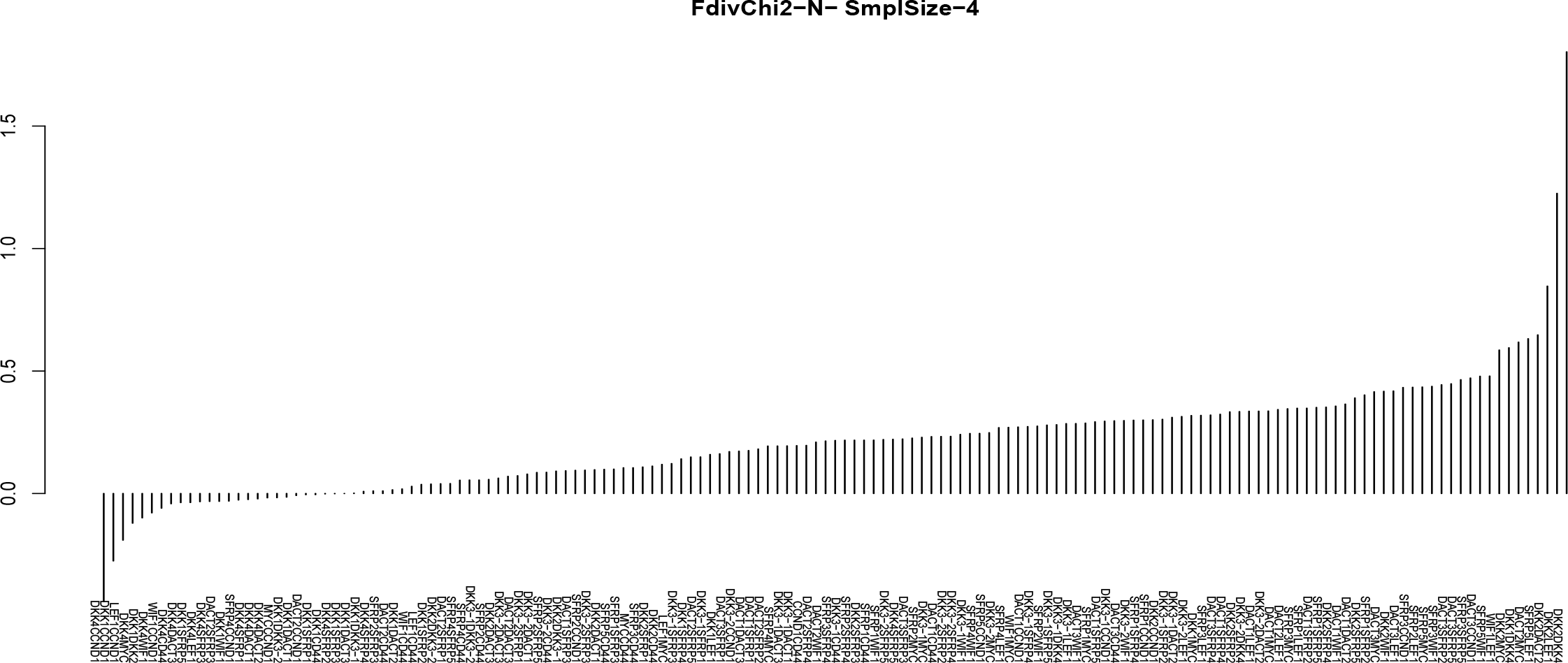
FdivChi2; Training sample size - 4; Test sample size - 16; Case - Normal

**Fig. 8.**
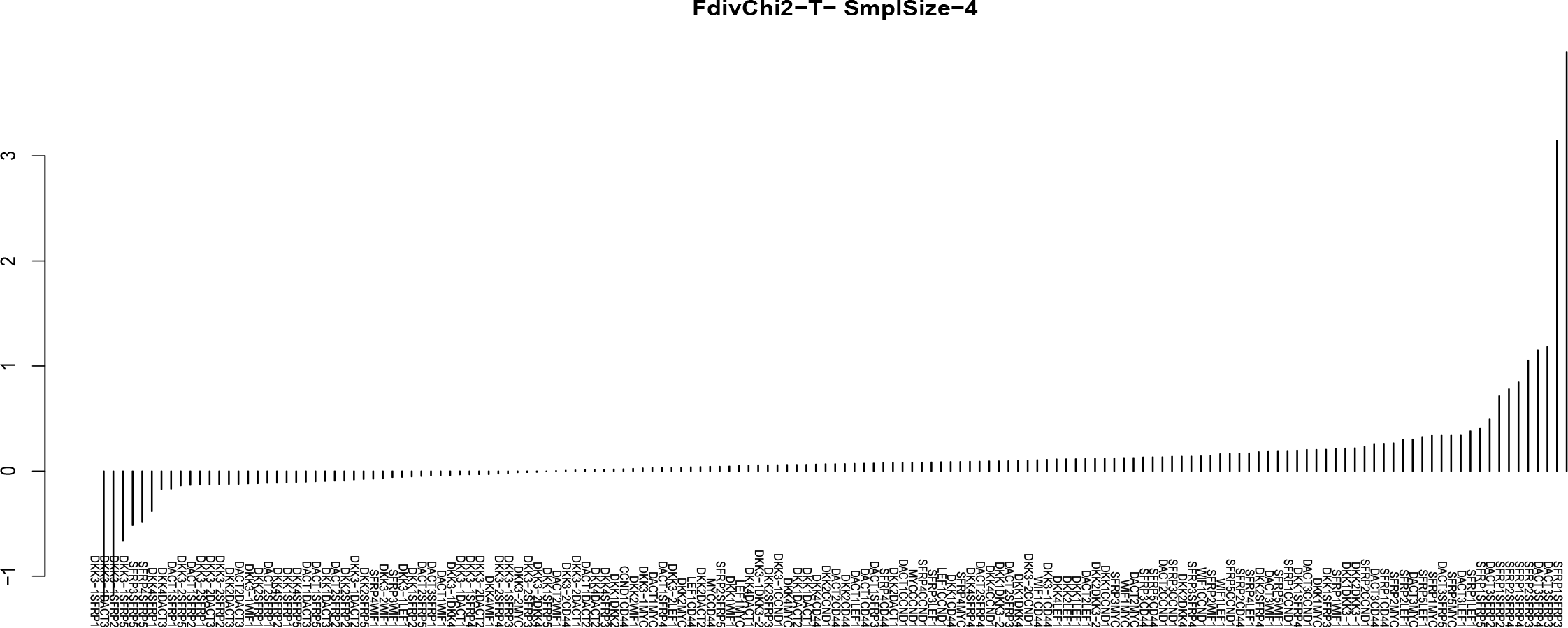
FdivChi2; Training sample size - 4; Test sample size - 16; Case - Tumor

**Fig. 9.**
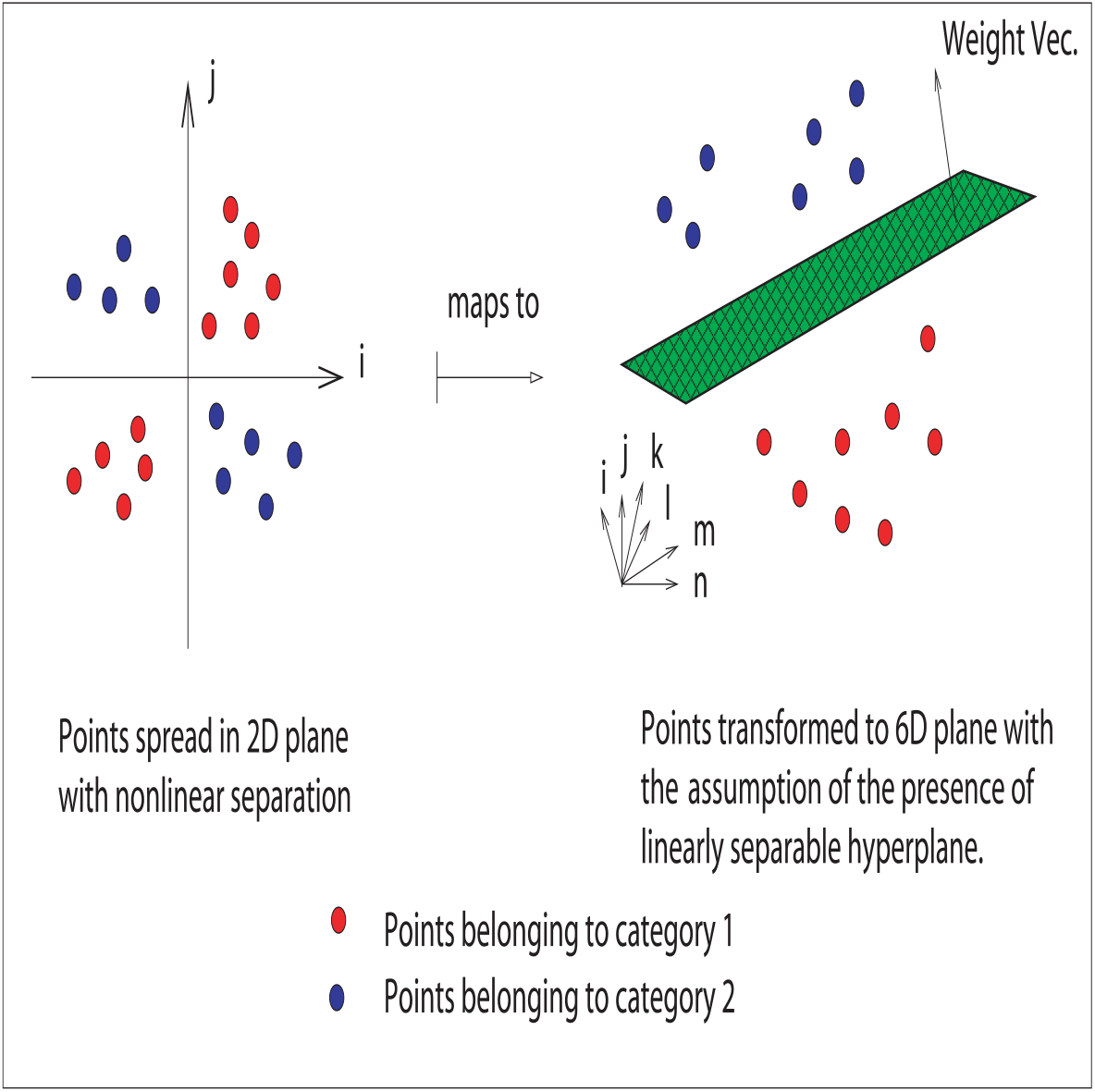
A geometrical interpretation of mapping nonlinearly separable data into higher dimensional space where it is assumed to be linearly separable, subject to the holding of dot product.

**Table 1.** Ranking of second order interactions in Tumor case using density and variance based sensitivity indices. Here 1 has high priority and 153 has low priority.

| Tumor Case - Density based methods |  |  |  |  |  |  |
| --- | --- | --- | --- | --- | --- | --- |
| Rank | Fdiv |  |  |  | HSIC |  |
| High priority interactions |  |  |  |  |  |  |
|  | chi2 | hellinger | kl | tv | rbf | laplace |
| 10 | SFRP1LEF1 | SFRP1MYC | SFRP5LEF1 | SFRP5LEF1 | SFRP2SFRP4 | SFRP1MYC |
| 9 | SFRP1SFRP5 | DACT3LEF1 | DACT3SFRP5 | DACT3SFRP5 | SFRP2SFRP3 | SFRP5LEF1 |
| 8 | DACT3SFRP2 | SFRP1SFRP2 | SFRP5MYC | SFRP5MYC | DACT3LEF1 | DACT3MYC |
| 7 | SFRP1SFRP2 | SFRP1LEF1 | SFRP1MYC | SFRP1MYC | SFRP1LEF1 | SFRP2LEF1 |
| 6 | SFRP2SFRP4 | SFRP2SFRP4 | DACT3SFRP4 | DACT3SFRP4 | SFRP1SFRP4 | DACT3LEF1 |
| 5 | SFRP1SFRP4 | SFRP2SFRP3 | DACT3MYC | DACT3MYC | DACT3SFRP4 | SFRP1WIF1 |
| 4 | SFRP2SFRP3 | SFRP1SFRP4 | DACT3LEF1 | DACT3LEF1 | SFRP1SFRP2 | DACT3WIF1 |
| 3 | DACT3SFRP4 | DACT3SFRP4 | SFRP1LEF1 | SFRP1LEF1 | DACT3SFRP2 | DACT3SFRP3 |
| 2 | DACT3SFRP3 | DACT3SFRP3 | DACT3SFRP3 | DACT3SFRP3 | DACT3SFRP3 | SFRP1LEF1 |
| 1 | SFRP1SFRP3 | SFRP1SFRP3 | SFRP1SFRP3 | SFRP1SFRP3 | SFRP1SFRP3 | SFRP1SFRP3 |
| Low priority interactions |  |  |  |  |  |  |
| 153 | DKK3-1SFRP1 | DKK3-1SFRP1 | DKK3-1SFRP1 | DKK3-1SFRP1 | DKK4DACT3 | DKK4DACT3 |
| 152 | DKK3-1DACT3 | DKK3-1DACT3 | DKK3-1DACT3 | DKK3-1DACT3 | DACT2DACT3 | DKK3-1SFRP5 |
| 151 | DKK3-1SFRP2 | DKK3-1SFRP2 | DKK3-1SFRP2 | DKK3-1SFRP2 | DKK3-1SFRP5 | DKK4SFRP1 |
| 150 | DKK3-1SFRP5 | DKK3-1SFRP5 | DKK3-1SFRP5 | DKK3-1SFRP5 | DKK1DACT3 | DACT2DACT3 |
| 149 | SFRP3SFRP5 | SFRP3SFRP5 | SFRP3SFRP5 | SFRP3SFRP5 | DKK1SFRP1 | DKK3-1SFRP2 |
| 148 | SFRP4SFRP5 | SFRP4SFRP5 | DACT1SFRP1 | DACT1SFRP1 | DACT1SFRP1 | DACT2SFRP1 |
| 147 | DKK4SFRP1 | DACT1SFRP1 | SFRP4SFRP5 | SFRP4SFRP5 | DKK3-2DACT3 | DACT2SFRP1 |
| 146 | DKK4DACT3 | DACT1SFRP2 | DACT1DACT3 | DACT1DACT3 | DKK2SFRP1 | DKK4SFRP2 |
| 145 | DACT1SFRP1 | DACT1DACT3 | DACT1SFRP5 | DACT1SFRP5 | DACT2SFRP1 | DACT1SFRP1 |
| 144 | DKK3-2SFRP5 | DKK4SFRP1 | DKK2DACT3 | DKK2DACT3 | DKK4SFRP1 | DKK2SFRP1 |

**Table 2.** Ranking of second order interactions in Normal case using density and variance based sensitivity indices. Here 1 has high priority and 153 has low priority.

| Normal Case - Density based methods |  |  |  |  |  |  |
| --- | --- | --- | --- | --- | --- | --- |
| Rank | Fdiv |  |  |  | HSIC |  |
| High priority interactions |  |  |  |  |  |  |
|  | chi2 | hellinger | kl | tv | rbf | laplace |
| 10 | DACT3CCND1 | DKK2WIF1 | DKK2DKK4 | DACT3MYC | DKK2SFRP5 | DACT3SFRP1 |
| 9 | SFRP5WIF1 | DKK2SFRP5 | DKK2LEF1 | DKK2WIF1 | DKK2WIF1 | DACT3CCND1 |
| 8 | WIF1LEF1 | DACT3SFRP2 | DKK2DACT2 | DACT3SFRP2 | SFRP5LEF1 | DKK2DACT2 |
| 7 | DKK2MYC | SFRP5LEF1 | DKK2MYC | SFRP5LEF1 | DACT3SFRP5 | DKK4SFRP5 |
| 6 | DKK1DKK4 | DACT3CCND1 | DACT3LEF1 | DACT3CCND1 | DACT3CCND1 | SFRP5CCND1 |
| 5 | DACT2MYC | DKK1MYC | DACT3CCND1 | DACT3LEF1 | SFRP5CCND1 | DACT3SFRP5 |
| 4 | SFRP5LEF1 | DKK2MYC | SFRP5LEF1 | DKK2MYC | DKK2LEF1 | DKK2SFRP5 |
| 3 | DKK2DACT2 | DKK2DACT2 | DACT3SFRP2 | DKK2DACT2 | DKK1MYC | DKK3-1CD44 |
| 2 | DKK2LEF1 | DKK2LEF1 | DKK2WIF1 | DKK2LEF1 | DKK2DACT2 | DKK2DKK4 |
| 1 | DKK2DKK4 | DKK2DKK4 | DACT3MYC | DKK2DKK4 | DKK2DKK4 | DKK1MYC |
| Low priority interactions |  |  |  |  |  |  |
| 153 | DKK4CCND1 | DKK4LEF1 | DKK4CD44 | DKK4CD44 | DKK1DKK3-2 | DKK4LEF1 |
| 152 | DKK1CCND1 | DKK1DKK3-2 | DKK4DACT3 | DKK4DACT3 | DKK4SFRP3 | DKK1DKK3-2 |
| 151 | LEF1CCND1 | DKK4SFRP1 | DKK4MYC | DKK4MYC | DKK4SFRP1 | DKK1WIF1 |
| 150 | DKK4MYC | DKK4SFRP3 | DKK4SFRP3 | DKK4SFRP3 | DKK4LEF1 | DKK4SFRP1 |
| 149 | DKK1DKK2 | DKK4DACT1 | DKK1DKK2 | DKK1DKK2 | DKK1SFRP3 | DKK3-2CD44 |
| 148 | DKK4WIF1 | DKK4DACT3 | DKK4DACT1 | DKK4DACT1 | DACT2SFRP3 | DKK3-2SFRP3 |
| 147 | WIF1CCND1 | DKK4CD44 | DKK4SFRP1 | DKK4SFRP1 | DKK4DACT1 | DKK3-2DACT3 |
| 146 | DKK4CD44 | DKK1SFRP1 | LEF1CCND1 | LEF1CCND1 | DKK1WIF1 | DKK3-2DACT1 |
| 145 | DKK4DACT3 | DKK1SFRP3 | DKK1DKK3-2 | DKK1DKK3-2 | DKK1SFRP1 | DKK3-2SFRP4 |
| 144 | DKK1SFRP5 | DACT2SFRP3 | DACT2SFRP3 | DACT2SFRP3 | WIF1CD44 | DKK3-2SFRP1 |

**Table 3.** Ranking of third order interactions in Tumor case using variance based sensitivity indices. Here 1 has high priority and 816 has low priority.

| Ranking | Tumor Case |  |  |
| --- | --- | --- | --- |
|  | HSIC |  |  |
|  | rbf | laplace | linear |
| High priority interactions |  |  |  |
| 10 | SFRP1SFRP2LEF1 | DACT3SFRP1WIF1 | DDK3-1DACT1SFRP4 |
| 9 | SFRP1SFRP2SFRP4 | DACT3SFRP1MYC | DDK3-1SFRP2SFRP3 |
| 8 | DACT3SFRP2LEF1 | DACT3SFRP5LEF1 | DACT3SFRP1SFRP4 |
| 7 | SFRP1SFRP2SFRP3 | DACT3SFRP2LEF1 | DDK3-1SFRP1SFRP3 |
| 6 | DACT3SFRP2SFRP4 | DACT3SFRP1SFRP2 | DDK3-1DACT3SFRP3 |
| 5 | DACT3SFRP1LEF1 | DACT3SFRP2SFRP4 | DACT3SFRP2SFRP4 |
| 4 | DACT3SFRP2SFRP3 | SFRP1SFRP2SFRP4 | DDK3-1SFRP2SFRP4 |
| 3 | DACT3SFRP1SFRP4 | DACT3SFRP1SFRP4 | DACT3SFRP1SFRP2 |
| 2 | DACT3SFRP1SFRP2 | DACT3SFRP1LEF1 | DDK3-1SFRP1SFRP4 |
| 1 | DACT3SFRP1SFRP3 | DACT3SFRP1SFRP3 | DDK3-1DACT3SFRP4 |
| Low priority interactions |  |  |  |
| 816 | DDK1DDK3-1SFRP2 | SFRP1SFRP3SFRP5 | DDK1DDK4WIF1 |
| 815 | DACT3SFRP3SFRP5 | DACT3SFRP4SFRP5 | DDK1DDK2DACT3 |
| 814 | SFRP1SFRP3SFRP5 | SFRP2SFRP4SFRP5 | DDK2DACT2WIF1 |
| 813 | DDK1DDK4SFRP1 | SFRP1SFRP4SFRP5 | DDK1DDK4SFRP1 |
| 812 | SFRP2SFRP3SFRP5 | SFRP2SFRP3SFRP5 | DDK2DDK4SFRP1 |
| 811 | DDK1DDK3-1SFRP5 | DDK1DDK3-1SFRP5 | DDK1DDK4LEF1 |
| 810 | DDK2DDK3-1SFRP5 | DDK1DDK3-1SFRP2 | DDK1DACT2LEF1 |
| 809 | DDK1DDK4DACT3 | DACT3SFRP3SFRP5 | DDK1DDK4SFRP2 |
| 808 | DDK2DDK4SFRP1 | DDK1DDK3-2SFRP1 | DDK1DDK2WIF1 |
| 807 | SFRP1SFRP4SFRP5 | DDK1DACT1SFRP2 | DACT3SFRP2WIF1 |

**Table 4.**
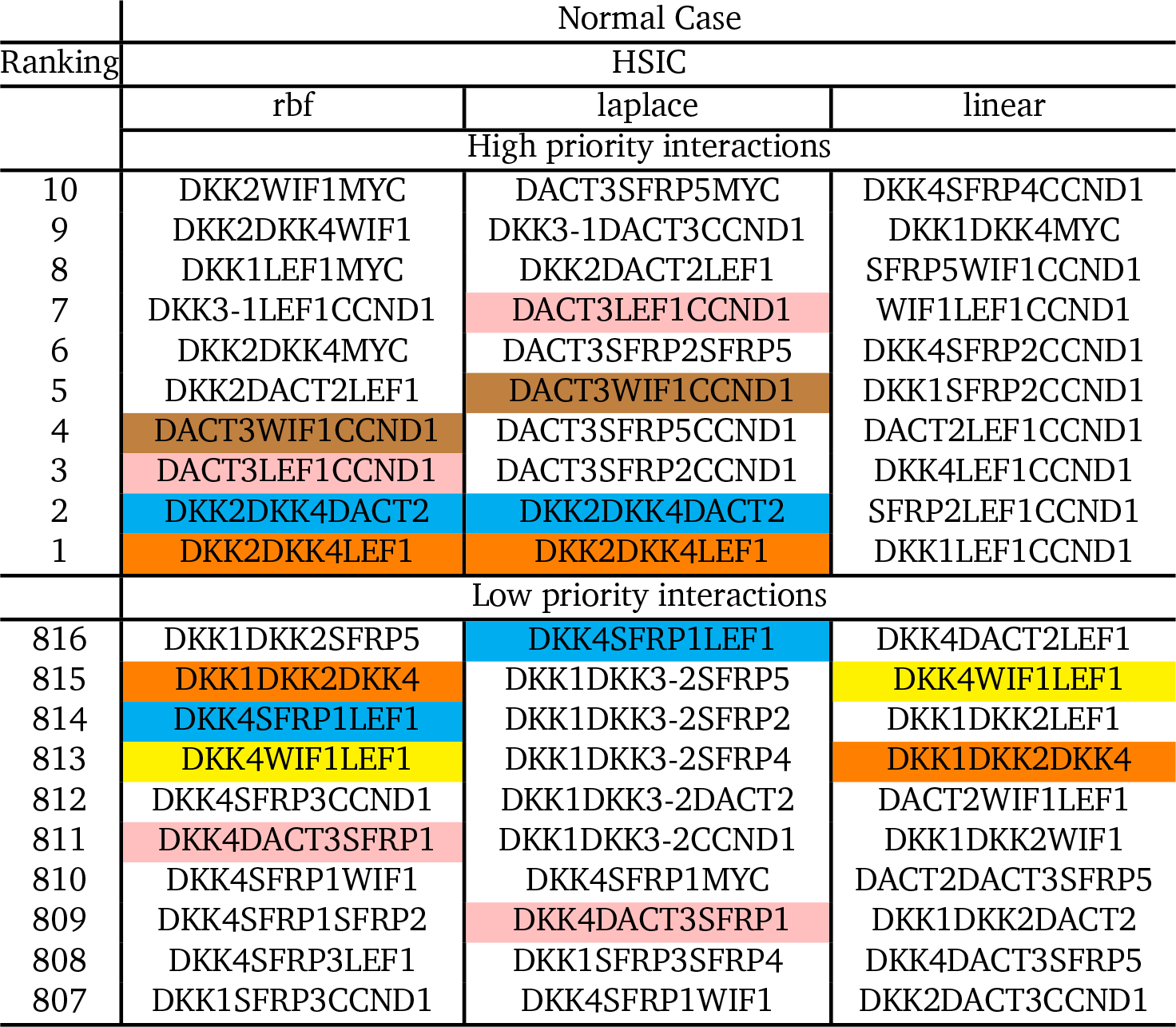
Ranking of third order interactions in Normal case using variance based sensitivity indices. Here 1 has high priority and 816 has low priority.

**Table 5.** Ranking of second order interactions in Tumor & Normal case using density and variance based sensitivity indices. Here 1 has high priority and 153 has low priority.

| Variance Based Method |  |  |  |  |  |  |
| --- | --- | --- | --- | --- | --- | --- |
|  | Tumor Case |  |  | Normal Case |  |  |
| High priority interactions |  |  |  |  |  |  |
| Rank | 2007 | jansen | martinez | 2007 | jansen | martinez |
| 10 | DACT3WIF1 | SFRP3LEF1 | SFRP3LEF1 | DKK3-2CD44 | SFRP3LEF1 | DKK3-1DACT2 |
| 9 | SFRP1LEF1 | DKK2LEF1 | DKK3-1MYC | SFRP2LEF1 | DKK2LEF1 | SFRP3WIF1 |
| 8 | DACT3LEF1 | DACT1LEF1 | DKK3-1DACT2 | DKK3-2WIF1 | DACT1LEF1 | SFRP3LEF1 |
| 7 | SFRP4LEF1 | DKK3-1WIF1 | DKK2LEF1 | DKK3-2DACT2 | DKK2DKK4 | DACT1LEF1 |
| 6 | DKK3-1WIF1 | DKK2DKK4 | DACT1LEF1 | WIF1MYC | SFRP4LEF1 | DKK2DKK4 |
| 5 | DACT1LEF1 | DKK3-2DKK4 | SFRP5LEF1 | DKK4MYC | DKK3-2DKK4 | SFRP5LEF1 |
| 4 | DKK3-2LEF1 | SFRP4LEF1 | DKK3-1LEF1 | DKK3-2DKK4 | DKK3-1DKK4 | DKK2LEF1 |
| 3 | SFRP4WIF1 | DKK3-2LEF1 | DKK3-1WIF1 | DKK3-2MYC | DKK3-1WIF1 | DKK3-1LEF1 |
| 2 | DACT2LEF1 | DKK3-1DKK4 | DKK2DKK4 | LEF1MYC | DKK3-2LEF1 | DKK3-1WIF1 |
| 1 | SFRP5LEF1 | SFRP5LEF1 | DKK3-1DKK4 | DKK3-2LEF1 | SFRP5LEF1 | DKK3-1DKK4 |
| Low priority interactions |  |  |  |  |  |  |
| 153 | DKK4MYC | LEF1CCND1 | DKK4SFRP3 | DKK1DKK4 | LEF1CCND1 | DKK4SFRP3 |
| 152 | DKK4DACT2 | DKK4MYC | DKK1DKK3-1 | DKK1LEF1 | DKK1SFRP5 | DKK1DKK3-1 |
| 151 | DKK1MYC | DKK1SFRP5 | DKK1SFRP3 | DKK4LEF1 | DKK4MYC | DKK4DACT1 |
| 150 | LEF1MYC | DKK4SFRP4 | DKK4DACT1 | DKK1WIF1 | DKK4SFRP3 | DKK1SFRP3 |
| 149 | DKK4CCND1 | DKK4SFRP3 | DKK4SFRP4 | WIF1CD44 | DKK4SFRP4 | DKK4SFRP4 |
| 148 | DKK1DACT2 | DKK4SFRP5 | DKK4CD44 | DKK1DKK3-2 | DKK1DKK3-2 | LEF1CCND1 |
| 147 | DKK4SFRP5 | DKK4DACT1 | DKK1DKK2 | DACT2LEF1 | DKK4SFRP5 | DKK4CD44 |
| 146 | DACT2MYC | DKK4CCND1 | DKK1DACT1 | DKK4SFRP2 | DKK1CCND1 | DKK1DKK2 |
| 145 | LEF1CCND1 | DKK1DKK3-2 | DKK4MYC | DKK1SFRP2 | DKK4DACT1 | DKK1DACT1 |
| 144 | DKK1CCND1 | DKK1CCND1 | LEF1CCND1 | DKK4CD44 | DKK4CCND1 | DKK1SFRP5 |

### 6.1 2*^nd^* Order interactions

In tumor cases (table 1), for most of the sensitvity indices we find that *SFRP*1-*SFRP*3 has the highest priority in the this data. This is followed by various others interactions like *DACT*3-*SFRP*3 and *DACT*3-*SFRP*4. For other kinds of the interactions we find varying kinds of rankings. It is important to note that the rankings are relative but they do give a glimpse of the importance of the interactions in a vast combinatorial space. These rankings are of importance in a particular phenomena of investigation. A particular colour across the columns, more specifically the sensitivity methods, represent the relative positioning of a particular interaction. The different colours in a row of a column do not mean any thing. What has been found is that *DACT*3 is implicated in the Wnt *β*-catenin signaling in colorectal cancer through epigenetic modifications and is a histone modification therapeutic target Jiang *et al*. ^16^. As a confirmatory result, the pipeline indicates the priority that *DACT*3 gets along with the members of the *SFRP* family in the tumor. This high priority also indicates the efficacy of the designed pipeline. Combinations of the members of the *SFRP* family also show at the high priority list. More specifically *SFRP*1-*SFRP*3, *SFRP*2-*SFRP*3 and *SFRP*1-*SFRP*2.

On the contrary, when we look at the set of low priority interactions, there is a range of combinations involving the family of *DKK* and members of *DACT*, *SFRP*. Surpurisingly, specific interactions of within family members also show up at this stage. This is specially seen for *DACT*1-*DACT*3, *DACT*2-*DACT*3, *SFRP*3-*SFRP*5 and *SFRP*4-*SFRP*5. These indicate the non-effectiveness of the 2^*nd*^ order interactions among a family of protein in the tumor case. Based on these rankings it might be possible to narrow down the search for important interactions in the vast combinatorial search space. We also generated the similar rankings for the interactions between the same set of factors in normal case. These are represented in table 2.

In comparison to the tables 1 and 2, which take into account the density based sensitivity methods, we also generated the rankings in tumor and normal cases using variance based sensitivity indices (table 5 in appendix). What we find is that there are interactions that get high priority in both the tumor and normal case. For example, the cases of *DKK*3 – 1-*WIF*1 and *DKK*3 – 2-*LEF*1, show up with high priority in both the normal and tumor case. This indicates the inability of the variance based sensitivity indices to capture crucial influence of the participating factors or combination of the factors, properly. Clearly, what we find is that the density based methods show superior selection and capture of improtant nonlinear biological information to provide proper indices that can be ranked later on.

### 6.2 3*^rd^* Order Interactions

We now turn our attention to the 3^*rd*^ order combinations which open a deeper and a greater space of investigation as compared to the space of the second order combinations. Tables 3 and 4 show the rankings for the 3^*rd*^ order interactions in tumor and normal cases, respectively. A major finding is that *DACT*3 combination with members of the family of *SFRP* show very high ranking in tumor cases. Note that we could only generate the rankings for the different kernels in the HSIC method as other density based methods were computationally intensive and costly in time. On the other hand the combinations of the family of *DKK* with other factors showed very low rankings in the tumor case, thus indicating their non participation or down regulated participation in the tumor cases. The most beautiful aspect of these rankings is that one can specifically see which members of a family are having potent role in the tumor and which ones are not. For example, *DACT*3 plays very important role along with various members of the *SFRP* family but the combination *DACT*3-*SFRP*3-*SFRP*5 shows very low participation in tumor case. This potential of the search engine to specifically rank the different combinations gives the biologists deep insights into the interactions of member of a single family of protiens also. A similar ranking profile was generated for the normal case.

CODE AVAILABILITY Code has been made available on Google drive at https://drive.google.com/folderview?id=0B7Kkv8wlhPU-V1Fkd1dMSTd5ak0&usp=sharing

## 7 Conclusions

A workflow has been presented that can prioritize the entire range of interactions among the constituent or subgroup of intra/extracellular factors affecting the pathway by using powerful algorithm of support vector ranking on interactions that have sensitivity indices of the involved factors as features. These sensitivity indices compute the influences of the factors on the pathway and represent nonlinear biological relations among the factors that are captured using kernel methods. SVM ranking then scores the testing data which can be sorted to find the highly prioritized interactions that need further investigation. Using this efficient workflow, it is possible to analyse any combination of involved factors in a signaling pathway. Here, we show confirmatory results for members of family of *DACT*, *SFRP* (*DKK*) to have highly prioritized role in tumor (normal) cases, in a single experimental setup. These preliminary results show the efficacy of the pipeline in providing a ranked list of interactions which the biologists can narrow down to in a vast serach space of combinations, thus cutting down the cost of terms of time, investment and energy. Additionally, the pipeline also generates unknown/untested biological hypothesis reagrding the combinations involved in the pathway in both tumor and normal cases. This opens up the potential for the biologists to work on recommendations from this pipeline in a vast combinatorial forest of combinations, when many a times, it is not possible to know where to start the search.

## Acknowledgements

### 8 Acknowledgement

The author thanks Mr. Prabhat Sinha and Mrs. Rita Sinha for financially supporting the project.

## Appendix

### 9 Sensitivity indices

#### 9.1 Variance based indices

The variance based indices as proposed by Sobol’ ^20^ prove a theorem that an integrable function can be decomposed into summands of different dimensions. Also, a Monte Carlo algorithm is used to estimate the sensitivity of a function apropos arbitrary group of variables. It is assumed that a model denoted by function *u* **=***ƒ* (*X*), *X* = (*X*_1_, *X*_2_, *…, X_n_*), is defined in a unit *n*-dimensional cube *K^n^* with *u* as the scalar output. The requirement of the problem is to find the sensitivity of function *ƒ* (*X*) with respect to different variables. If *u*^*^ **=** *ƒ* (*X*^*^) is the required solution, then the sensitivity of *u*^*^ apropos *X_k_* is estimated via the partial derivative (*∂u/ ∂X_k_*)_*X***=***X*_*. This approach is the local sensitivity. In global sensitivity, the input *X* **=** *X*^*^ is not specified. This implies that the model *ƒ* (*X*) lies inside the cube and the sensitivity indices are regarded as tools for studying the model instead of the solution. Detailed technical aspects with examples can be found in Homma and Saltelli ^32^ and **?**.

Let a group of indices *i*_1_, *i*_2_, *…, i_s_* exist, where 1 ≤ *i*_1_ *< … < i_s ≤_ n* and 1 ≤ *s ≤ n*. Then the notation for sum over all different groups of indices is -

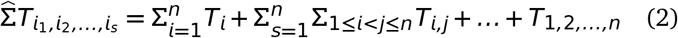

Then the representation of *ƒ* (*X*) using equation 2 in the form -

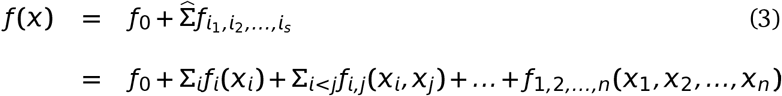

is called ANOVA-decomposition from Archer *et al*. ^45^ or expansion into summands of different dimensions, if *ƒ*_0_ is a constant and integrals of the summands *ƒ_i_*_1, *i*2, *…,is*_ with respect to their own variables are zero, i.e,

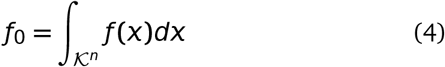

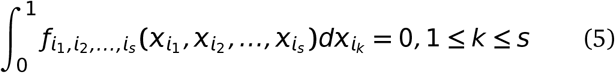

It follows from equation 4 that all summands on the right hand side are orthogonal, i.e if at least one of the indices in *i*_1_, *i*_2_, *…, i_s_* and *j*_1_, *j*_2_, *…, j_l_* is not repeated i.e

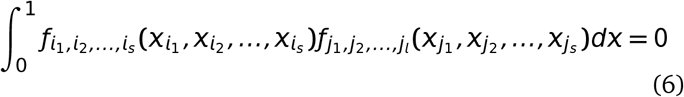

Sobol’ ^20^ proves a theorem stating that there is an existence of a unique expansion of equation 4 for any *ƒ* (*X*) integrable in *K^n^*. In brief, this implies that for each of the indices as well as a group of indices, integrating equation 4 yields the following –

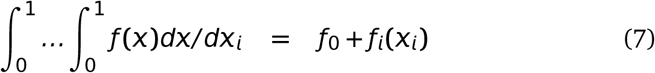

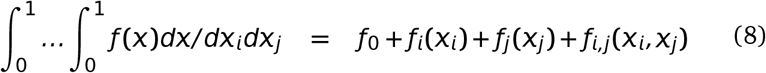

were, *dX/ dX_i_* is ∏_∀*k*∈{1,..,*n*};*i*∉ *k*_ dX_*k*_ and *dX/ dX_i_dX_j_* is ∏_∀*k*∈{1,..,*n*};*i*∉ *k*_ dX_k_. For higher orders of grouped indices, similar computations follow. The computation of any summand *ƒ_i_*_1_, *i*2, *…,i_s_* (*X_i_*_1_, *X_i_*_2_, *…, X_is_*) is reduced to an integral in the cube *K^n^*. The last summand *ƒ*_1,2,*…,n*_(*X*_1_, *X*_2_, *…, X_n_*) is *ƒ* (*X***) –** *ƒ*_0_ from equation 4. Homma and Saltelli ^32^ stresses that use of Sobol sensitivity indices does not require evaluation of any *ƒ_i_*_1, *i*2, *…,is*_ (*X_i_*_1_, *X_i_*_2_, *…, X_is_*) nor the knowledge of the form of *ƒ* (*X*) which might well be represented by a computational model i.e a function whose value is only obtained as the output of a computer program.

Finally, assuming that *ƒ* (*X*) is square integrable, i.e *ƒ* (*X*) ∈ *L*_2_, then all of *ƒ_i_*_1, *i*2, *…,is*_ (*X_i_*_1_, *X_i_*_2_, *…, X_is_*) *L*_2_. Then the following constants

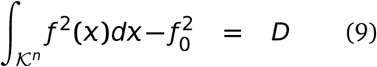

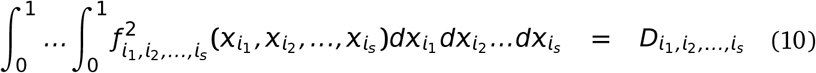

are termed as variances. Squaring equation 4, integrating over and using the orthogonality property in equation 6, *D* evaluates to -

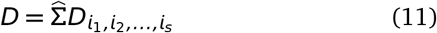

Then the global sensitivity estimates is defined as –

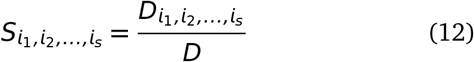

It follows from equations 11 and 12 that

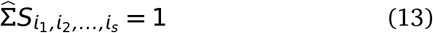

Clearly, all sensitivity indices are non-negative, i.e an index *S_i_*_1, *i*2, *…,is*_ = 0 if and only if *ƒ_i_*_1, *i*2, *…,is*_ 0. The true potential of Sobol indices is observed when variables *X*_1_, *X*_2_, *…, X_n_* are divided into *m* different groups with *y*_1_, *y*_2_, *…, y_m_* such that *m < n*. Then *ƒ* (*X*) *ƒ* (*y*_1_, *y*_2_, *…, y_m_*). All properties remain the same for the computation of sensitivity indices with the fact that integration with respect to *y_k_* means integration with respect to all the *X_i_*’s in *y_k_*. Details of these computations with examples can be found in **?** Variations and improvements over Sobol indices have already been stated in section 3.

#### 9.2 Density based indices

As discussed before, the issue with variance based methods is the high computational cost incurred due to the number of interactions among the variables. This further requires the use of screening methods to filter out redundant or unwanted factors that might not have significant impact on the output. Recent work by Da Veiga ^56^ proposes a new class of sensitivity indicies which are a special case of density based indicies Borgonovo ^53^. These indicies can handle multivariate variables easily and relies on density ratio estimation. Key points from Da Veiga ^56^ are mentioned below.

Considering the similar notation in previous section, *ƒ*: *R^n^* →*R* (*u* **=** *ƒ* (*X*)) is assumed to be continuous. It is also assumed that *X_k_* has a known distribution and are independent. Baucells and Borgonovo ^68^ state that a function which measures the similarity between the distribution of *U* and that of *U X_k_* can define the impact of *X_k_* on *U*. Thus the impact is defined as –

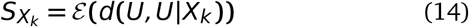

were *d*(⋅, ⋅) is a dissimilarity measure between two random variables. Here *d* can take various forms as long as it satisfies the criteria of a dissimilarity measure. Csiszár *et al*. ^59^’s f-divergence between *U* and *U |X_k_* when all input random variables are considered to be absolutely continuous with respect to Lebesgue measure on *R* is formulated as –

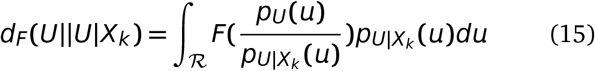

were *F* is a convex function such that *F*(1**) =** 0 and *p_U_* and *pU |X_k_* are the probability distribution functions of *U* and *U| X_k_*. Standard choices of *F* include Kullback-Leibler divergence *F*(*t***) =–**log_*e*_(*t*), Hellinger distance 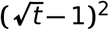, Total variation distance *F*(*t***) =** |*t* − 1|, Pearson *χ*^2^ divergence *F*(*t***) =** *t*^2^ − 1 and Neyman *χ*^2^ divergence *F*(*t*) = (1 − *t*^2^)*/ t*. Substituting equation 15 in equation 14, gives the following sensitivity index -

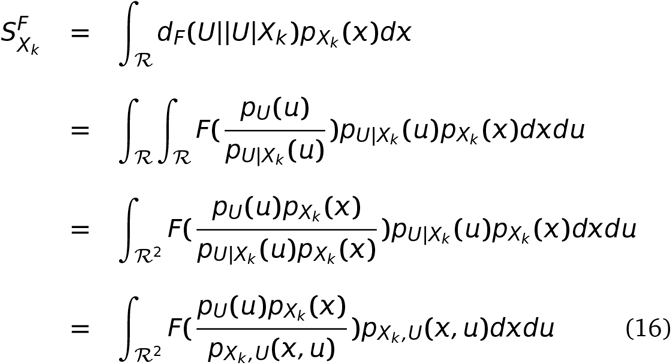

were *p X_k_* and *pX_k_,Y* are the probability distribution functions of *X_k_* and (*X_k_, U*), respectively. Csiszár *et al*. ^59^ f-divergences imply that these indices are positive and equate to 0 when *U* and *X_k_* are independent. Also, given the formulation of 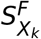, it is invariant under any smooth and uniquely invertible transformation of the variables *X_k_* and *U* (Kraskov *et al*. 69). This has an advantage over Sobol sensitivity indices which are invariant under linear transformations.

By substituting the different formulations of *F* in equation 16, Da Veiga ^56^’s work claims to be the first in establishing the link that previously proposed sensitivity indices are actually special cases of more general indices defined through Csiszár *et al*. ^59^’s f-divergence. Then equation 16 changes to estimation of ratio between the joint density of (*X_k_, U*) and the marginals, i.e -

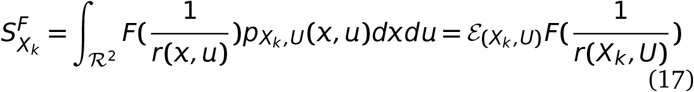

were, *r*(*X, y*) = (*p_Xk,U_* (*X, u*))*/* (*p_U_* (*u*)*p_Xk_* (*X*)). Multivariate extensions of the same are also possible under the same formulation.

Finally, given two random vectors *X* ∈ *𝓡^p^* and *Y* ∈*𝓡^q^*, the dependence measure quantifies the dependence between with the property that the measure equates to 0 if and only if *X* and *Y* are independent. These measures carry deep links (Sejdinovic *et al*. ^70^) with distances between embeddings of distributions to reproducing kernel Hilbert spaces (RHKS) and here the related Hilbert-Schmidt independence criterion (HSIC by Gretton *et al*. ^58^) is explained.

In a very brief manner from an extremely simple introduction by Daumé III ^71^ -”We first defined a field, which is a space that supports the usual operations of addition, subtraction, multiplication and division. We imposed an ordering on the field and described what it means for a field to be complete. We then defined vector spaces over fields, which are spaces that interact in a friendly way with their associated fields. We defined complete vector spaces and extended them to Banach spaces by adding a norm. Banach spaces were then extended to Hilbert spaces with the addition of a dot product.” Mathematically, a Hilbert space *ℌ* with elements *r, s* ∈ *ℌ* has dot product 〈*r, s*〉_*ℌ*_ and *r* · *s*. When *ℌ* is a vector space over a field *𝓕*, then the dot product is an element in *𝓕*. The product 〈*r, s*〉_*ℌ*_ follows the below mentioned properties when *r, s, t* ∈ *ℌ* and for all *a* ∈ *𝓕* -

- Associative: (*ar*) · *s* = *a*(*r* · *s*)
- Commutative: *r* · *s* = *s* · *r*
- Distributive: *r* · (*s* **+** *t*) = *r* · *s* **+** *r* · *t*

Given a complete vector space *V* with a dot product 〈·, ·〉, the norm on *V* defined by ||*r*|| *_V_* = 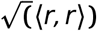 makes this space into a Banach space and therefore into a full Hilbert space.

A reproducing kernel Hilbert space (RKHS) builds on a Hilbert space *ℌ* and requires all Dirac evaluation functionals in *ℌ* are bounded and continuous (on implies the other). Assuming *ℌ* is the *L*_2_ space of functions from *X* to *R* for some measurable, *X* is a vector and *g* a function which maps from this *X*. For an element *X* ∈ *X*, a Dirac evaluation functional at x is a functional *δ_X_* ∈*ℌ* such that *δ_X_*(*g***) =** *g*(*X*). For the case of real numbers, X. is a vector and *g* a function which maps from this vector space to *R*. Then *δ_X_* is simply a function which maps *g* to the value *g* has at *X*. Thus, *δ_X_* is a function from (*𝓡^n^* → *𝓡*) into *𝓡*.

The requirement of Dirac evaluation functions basically means (via the Riesz ^72^ representation theorem) if *ϕ* is a bounded linear functional (conditions satisfied by the Dirac evaluation functionals) on a Hilbert space *ℌ*, then there is a unique vector *ℓ* in *ℌ* such that *ϕg* =〈 *g, ℓ* 〉_*ℌ*_ for all *ℓ*∈ *H*. Translating this theorem back into Dirac evaluation functionals, for each *δ_X_* there is a unique vector *k_X_* in *ℌ* such that *δ_X_g* = *g*(*X*) = 〈 *g, k_X_* 〉 _*H*_. The reproducing kernel *K* for *ℌ* is then defined as: *K*(*X, X*′) = 〈 *k_X_, k_X_*〉, were *k_X_* and *k_X_*′ are unique representatives of *δ_X_* and *δ_X_*′. The main property of interest is 〈*g*, *K*(*X, X*^I^) 〉 _*H*_ = *g* (*X* Furthermore, *k_x_* is defined to be a function *y* → *K*(*X, y*) and thus the reproducibility is given by 〈*K*(*X*, ·), *K*(*y*, ·)〉_*ℌ*_ = *K*(*X, y*).

Basically, the distance measures between two vectors represent the degree of closeness among them. This degree of closeness is computed on the basis of the discriminative patterns inherent in the vectors. Since these patterns are used implicitly in the distance metric, a question that arises is, how to use these distance metric for decoding purposes?

The kernel formulation as proposed by Aizerman *et al*. ^60^, is a solution to our problem mentioned above. For simplicity, we consider the labels of examples as binary in nature. Let **x**_*i*_ ∈*𝓡^n^*, be the set of *n* feature values with corresponding category of the example label (*y_i_*) in data set *D*. Then the data points can be mapped to a higher dimensional space *ℌ* by the transformation *ϕ*:

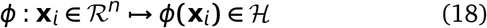

This *ℌ* is the *Hilbert Space* which is a strict inner product space, along with the property of completeness as well as separability. The inner product formulation of a space helps in discriminating the location of a data point w.r.t a separating hyperplane in *ℌ*. This is achieved by the evaluation of the inner product between the normal vector representing the hyperplane along with the vectorial representation of a data point in *ℌ*. Thus, the idea behind equation (18) is that even if the data points are nonlinearly clustered in space *R^n^*, the transformation spreads the data points into *ℌ*, such that they can be linearly separated in its range in *ℌ*.

Often, the evaluation of dot product in higher dimensional spaces is computationally expensive. To avoid incurring this cost, the concept of kernels in employed. The trick is to formulate kernel functions that depend on a pair of data points in the space *𝓡^n^*, under the assumption that its evaluation is equivalent to a dot product in the higher dimensional space. This is given as:

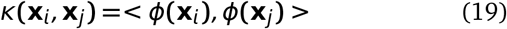

Two advantages become immediately apparent. First, the evaluation of such kernel functions in lower dimensional space is computationally less expensive than evaluating the dot product in higher dimensional space. Secondly, it relieves the burden of searching an appropriate transformation that may map the data points in *𝓡^n^* to *ℌ*. Instead, all computations regarding discrimination of location of data points in higher dimensional space involves evaluation of the kernel functions in lower dimension. The matrix containing these kernel evaluations is referred to as the *kernel* matrix. With a cell in the kernel matrix containing a kernel evaluation between a pair of data points, the kernel matrix is square in nature.

As an example in practical applications, once the kernel has been computed, a pattern analysis algorithm uses the kernel function to evaluate and predict the nature of the new example using the general formula:

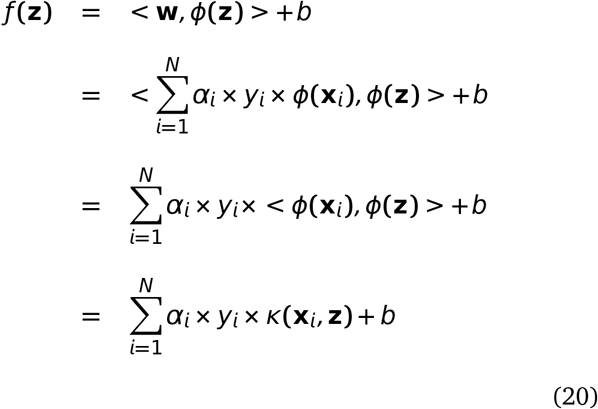

where **w** defines the hyperplane as some linear combination of training basis vectors, **z** is the test data point, *y_i_* the class label for training point **x**_*i*_, *α_i_* and *b* are the constants. Various transformations to the kernel function can be employed, based on the properties a kernel must satisfy. Interested readers are referred to Taylor and Cristianini ^73^ for description of these properties in detail.

The Hilbert-Schmidt independence criterion (HSIC) proposed by Gretton *et al*. ^58^ is based on kernel approach for finding dependences and on cross-covariance operators in RKHS. Let *X* ∈ *X* have a distribution *P_X_* and consider a RKHS *A* of functions *X* → *𝓡* with kernel *k_X_* and dot product 〈·, ·〉_*A*_. Similarly, Let *U* ∈ *Y* have a distribution *P_Y_* and consider a RKHS *B* of functions *U* →*𝓡* with kernel *k_B_* and dot product, 〈·, ·〉_*B*_. Then the cross-covariance operator *C_X,U_* associated with the joint distribution *P_XU_* of (*X, U*) is the linear operator *B* → *A* defined for every *a* ∈ *A* and *b* ∈ *B* as -

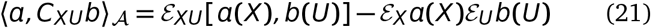

The cross-covariance operator generalizes the covariance matrix by representing higher order correlations between *X* and *U* through nonlinear kernels. For every linear operator *C*: *B*→ *A* and provided the sum converges, the Hilbert-Schmidt norm of *C* is given by -

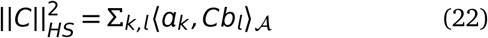

were *a_k_* and *b_l_* are orthonormal bases of *A* and *B*, respectively. The HSIC criterion is then defined as the Hilbert-Schmidt norm of cross-covariance operator –

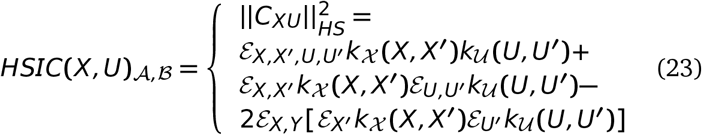

were the equality in terms of kernels is proved in Gretton *et al*. ^58^. Finally, assuming (*X_i_, U_i_*) (*i* **=** 1, 2, *…, n*) is a sample of the random vector (*X, U*) and denote *K_X_* and *K_U_* the Gram matrices with entries *K_X_* (*i, j*) = *k_X_* (*X_i_, X_j_*) and *K_U_* (*i, j*) = *k_U_* (*U_i_, U_j_*). Gretton *et al*. ^58^ proposes the following estimator for *HSIC_n_*(*X, U*)_*A,B*_ –

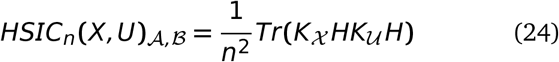

were *ℌ* is the centering matrix such that *ℌ*(*i,j*) = 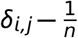. Then *HSIC_n_*(*X, U*)_*A,B*_ can be expressed as -

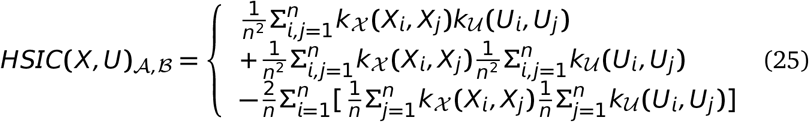

Finally, Da Veiga ^56^ proposes the sensitivity index based on distance correlation as -

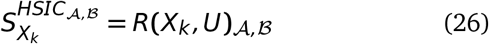

were the kernel based distance correlation is given by –

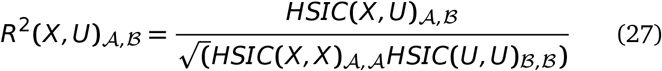

were kernels inducing *A* and *B* are to be chosen within a universal class of kernels. Similar multivariate formulation for equation 24 are possible.

#### 9.3 Choice of sensitivity indices

The SENSITIVITY PACKAGE (Faivre *et al*. ^74^ and Iooss and Lemaître ^21^) in R langauge provides a range of functions to compute the indices and the following indices will be taken into account for addressing the posed questions in this manuscript.

1. sensiFdiv - conducts a density-based sensitivity analysis where the impact of an input variable is defined in terms of dissimilarity between the original output density function and the output density function when the input variable is fixed. The dissimilarity between density functions is measured with Csiszar f-divergences. Estimation is performed through kernel density estimation and the function kde of the package ks. (Borgonovo ^53^, Da Veiga ^56^)
2. sensiHSIC - conducts a sensitivity analysis where the impact of an input variable is defined in terms of the distance between the input/output joint probability distribution and the product of their marginals when they are embedded in a Reproducing Kernel Hilbert Space (RKHS). This distance corresponds to HSIC proposed by Gretton *et al*. ^58^ and serves as a dependence measure between random variables.
3. soboljansen - implements the Monte Carlo estimation of the Sobol indices for both first-order and total indices at the same time (all together 2p indices), at a total cost of (p+2) × n model evaluations. These are called the Jansen estimators. (Jansen ^48^ and Saltelli *et al*. ^40^)
4. sobol2002 - implements the Monte Carlo estimation of the Sobol indices for both first-order and total indices at the same time (all together 2p indices), at a total cost of (p+2) × n model evaluations. These are called the Saltelli estimators. This estimator suffers from a conditioning problem when estimating the variances behind the indices computations. This can seriously affect the Sobol indices estimates in case of largely non-centered output. To avoid this effect, you have to center the model output before applying “sobol2002”. Functions “soboljansen” and “sobolmartinez” do not suffer from this problem. (Saltelli ^34^)
5. sobol2007 - implements the Monte Carlo estimation of the Sobol indices for both first-order and total indices at the same time (all together 2p indices), at a total cost of (p+2) × n model evaluations. These are called the Mauntz estimators. (Saltelli and Annoni ^47^)
6. sobolmartinez - implements the Monte Carlo estimation of the Sobol indices for both first-order and total indices using correlation coefficients-based formulas, at a total cost of (p+ 2) × n model evaluations. These are called the Martinez estimators.
7. sobol - implements the Monte Carlo estimation of the Sobol sensitivity indices. Allows the estimation of the indices of the variance decomposition up to a given order, at a total cost of (N + 1) × n where N is the number of indices to estimate. (Sobol’ ^20^)

### 10 Optimization and Support Vector Machines

Aspects of SVMs from Sinha ^75^ are reproduced for completeness.

#### 10.1 Optimization Problems

##### 10.1.1 Introduction

The main focus in this section is optimization problems, the concept of Lagrange multipliers and KKT conditions, which will be later used to explain the details about the SVMs.

##### 10.1.2 Mathematical Formulation

Optimization problems arise in almost every area of engineering. The goal is to achieve an almost perfect and efficient result, while carrying out certain procedures of optimization. Our main source of reference on this topic derives from Bletzinger ^76^. We will be using notations used in Bletzinger ^76^. In mathematical terms the general form of optimization problem can be represented as:

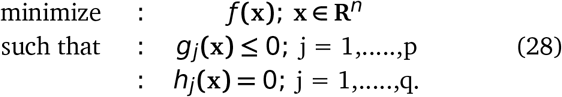

where *ƒ, g_j_* and *h_j_* are the objective function, equality constraints and inequality constraints. Generally, the number of constraints is less than the number of variables used to formulate the optimization problem. For a problem to be linear, both the constraints and the objective function need to be linear. Quadratic problems require only the objective function to be quadratic, while the constraints remain linear in formulation. Besides these, if any one of the functions is nonlinear, then the problem becomes nonlinear in nature. A graphical view of the types of the problems can be seen in fig. 10).

**Fig. 10.**
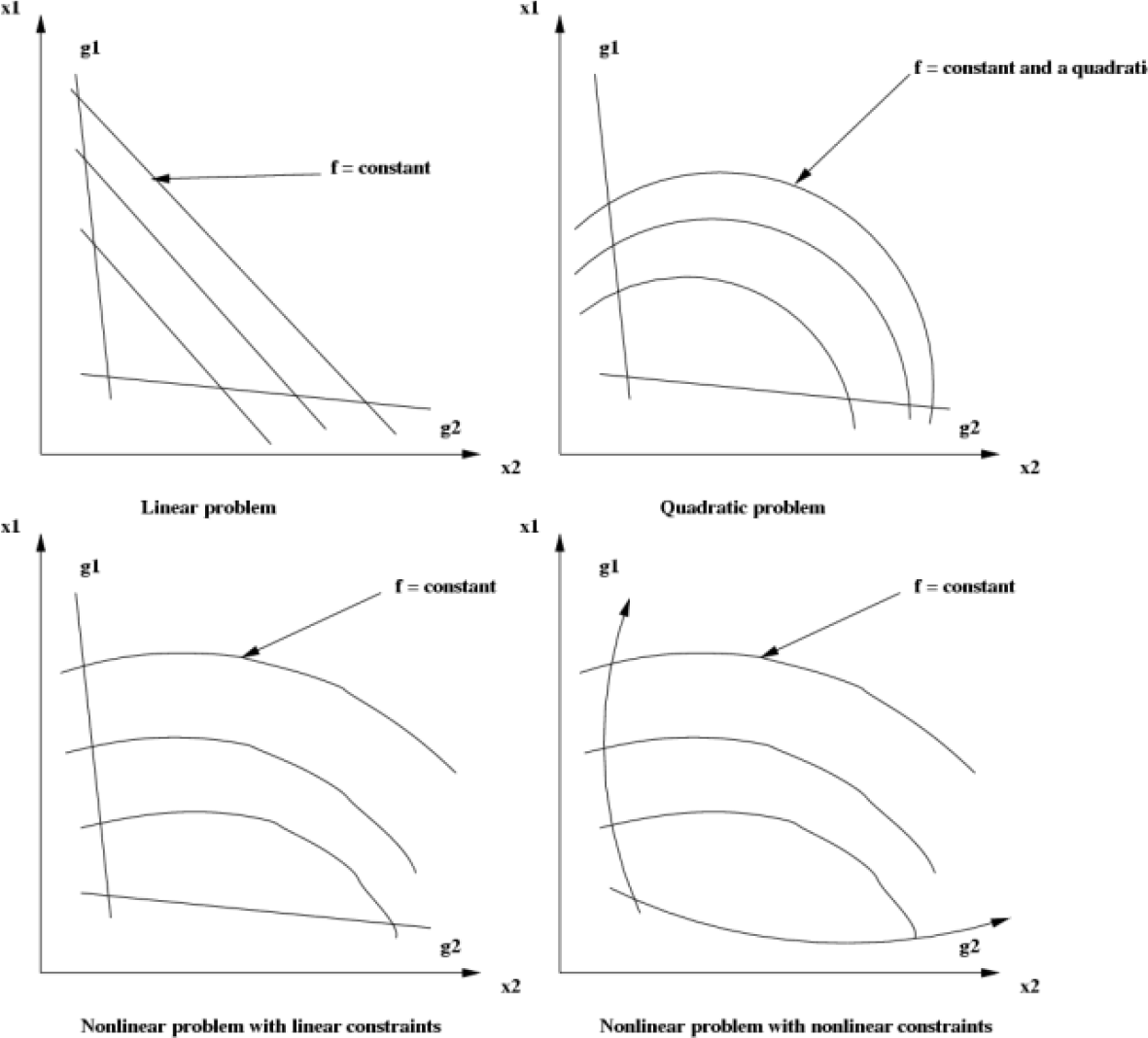
Kinds of optimization problems.

##### 10.1.3 Lagrange Multipliers

In unconstrained optimization problems, where the first order derivatives are assumed continuous, the solution is found by solving:

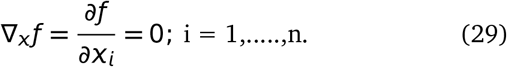

where *ƒ* is a function of *X*. Since most of the optimization problems are constrained, the concept of Lagrange multipliers is introduced in order to solve the problem. Thus, the Lagrangian formulation, for Eqn. 28 becomes:

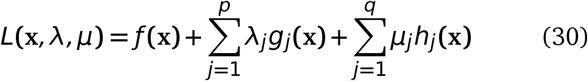

where *L* is the Lagrangian, *λ* and *µ* are the vectors of the Lagrange multipliers for inequality and equality constraints, respectively.

Next comes the solving of the Lagrangian. We try to derive a solution in terms of variables used and show that the final solution achieved by Equ. 28 and Eqn. 30 remains the same. For the sake of derivation, we assume that each of the vectors *X*, *λ* and *µ* have a single element and also there exists a single optimal solution. We will then generalize the solution to vectors containing various elements. Let *X*^*^, *λ*^*^ and *µ*^*^ be the optimal solution for the Lagrangian. Let *X*^!^ be the optimal solution for *ƒ* (*X*). To begin with, our Lagrangian has the form:

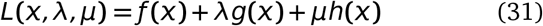

###### Derivation

- **Step 1:** Differentiate the Lagrangian in Eqn 31 w.r.t *X* and equate it to zero.

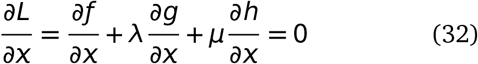
- **Step 2:** Find *X* in terms of *λ* and *µ*, such that *X* **=** *X*(*λ, µ*).
- **Step 3:** Differentiate the Lagrangian in Eqn. 31 w.r.t *λ* and equate it to zero.

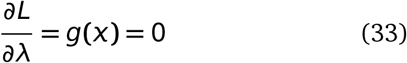
- **Step 4:** Differentiate the Lagrangian in Eqn 31 w.r.t *µ* and equate it to zero.

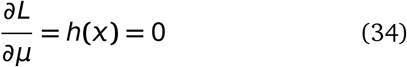
- **Step 5:** Substitute *X*(*λ, µ*) in Equ. 33 and Eqn. 34 to get two equations in two unknowns *λ* and *µ* and solve to get the optimal values.

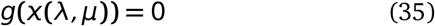

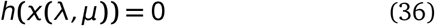 Let *λ*^*^, *µ*^*^ be the solution. Substituting these in *X* **=** *X*(*λ, µ*), we get *X*^*^.
- **Step 6:** Combining Eqn. 33 and Eqn. 34 in Eqn. 31, along with *λ*^*^, *µ*^*^ and *X*^*^, we have:

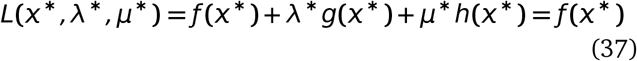 Since, it is assumed that there exist only one optimal solution we have:

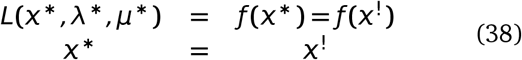

Lastly, since *g*(*X*) in Eqn. 33 is a inequality constraint, we have:

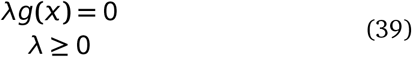

##### 10.1.4 Dual Functions

For sake of simplicity, let us for a moment ignore the equality constraint. Then the Lagrangian becomes:

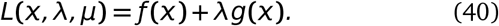

It is sometimes easy to transform the Lagrangian into a simpler form, in order to find an optimal solution. We can represent the Lagrangian as a Dual function in such a manner that the optimal solution defined as minimum of *L*(*X, λ*^*^) w.r.t *X* where *λ* **=** *λ*^*^, can be represented as the maximum of dual function *D*(*λ*) w.r.t *λ*. For a given *λ*, the dual is evaluated by finding the minimum of *L*(*X, λ*) w.r.t *X*. Thus to find the optimal point we evaluate:

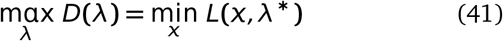

So the basic steps to solve the dual problem are as follows: **Step 1:** Minimize *L*(*X, λ*) w.r.t *X*, and find *X* in terms of *λ*. **Step 2:** Substitute *X*(*λ*) in *L* s.t. *D*(*λ***) =** *L*(*X*(*λ*), *λ*). **Step 3:** Maximize *D*(*λ*) w.r.t *λ*.

##### 10.1.5 Karush Kuhn Tucker Conditions

The derivation in the last part (Eqn. 31 to Eqn. 39)gives us a set of equations that need to be evaluated along with the consideration of constraints present. These set of equations and constraints in terms of the Lagrangian, form the Karush Kuhn Tucker Conditions. We give here the generalized KKT conditions and explain the necessary details

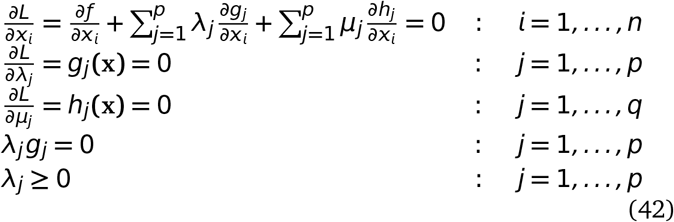

where *L* is Eqn. 30.

The KKT conditions specify a few points which are as follows:

1. The first line states that the linear combination of objective and constraint gradients vanishes.
2. A prerequisite of the KKT conditions is that the gradients of the constraints must be continuous (evident from second and third lines in Eqn. 42).
3. The last two lines in Eqn. 42 state that at optimum either the constraints are active or the constrains are inactive.

#### 10.2 Support Vector Machines

Armed with the knowledge of optimization problems and concept of Lagrange multipliers, we now delve into the workings of support vector machines. Burges ^77^ provides a good introduction to SVMs and is our main reference. Interested readers should refer to Cristianini and Shawe-Taylor ^78^, Schölkopf and Smola ^79^ and Vapnik and Vapnik ^80^ for detailed references.

##### 10.2.1 Separable Case

Let us suppose that we are presented with a data set that is linearly separable. We assume that there are *m* examples of data in the format {**x**_*i*_, *y_i_*}, s.t. **x**_*i*_ ∈ **R**^*n*^; *i* = 1, *…., m*, where *y_i_* ∈ {−1, 1} is the corresponding true label of **x**_*i*_. We also suppose there is an existence of a linear hyperplane in the *n* dimensional space that separates the positively labeled data from the negatively labeled data. Let this separating hyperplane be given by

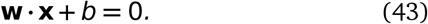

where, **w** is the normal vector to the hyperplane and |*b| / ‖***w**‖ is the shortest perpendicular distance of the hyperplane to the origin. **w** is the Euclidean norm of **w**. The *margin* of a hyperplane is then defined as the minimum of the distance of the positively and negatively labeled examples, to the hyperplane. For the linear case, the SVM searches for the hyperplane with largest margin. We now have three conditions, based on the location of an example **x**_*i*_ w.r.t the hyperplane:

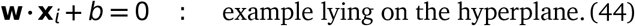

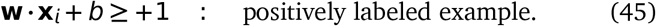

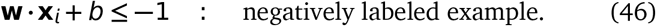

Combining the equality and the two inequalities we have:

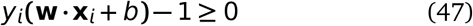

Since the SVMs search for the largest margin, we now try to find a mathematical expression of the margin. Considering the examples that satisfy equality in Eqn. 45, the distance of the closest positive example can be expressed as 1 − *b /* ||**w**||. Similarly, considering the negative examples that satisfy equality in Eqn. 46, the distance of the closest negative example can be expressed as | − 1 − *b| /*||**w**||. On summation of the two shortest distances, we get the margin of the hyperplane as 2*/* ||**w**||. Since the labels are { 1, 1}, no example lies inside the hyperplanes representing the margin in this case. Taking into account that the SVM searches for the largest margin, we can say that it can be achieved by minimizing ||**w**|| ^2^, subject to the constraints in Eqn. 47. Examples lying on the hyperplanes of the margins are termed *support vectors*, as their removal would change the margin and thus the solution. Figure 11 represents the conceptual points about separating hyperplanes.

**Fig. 11.**
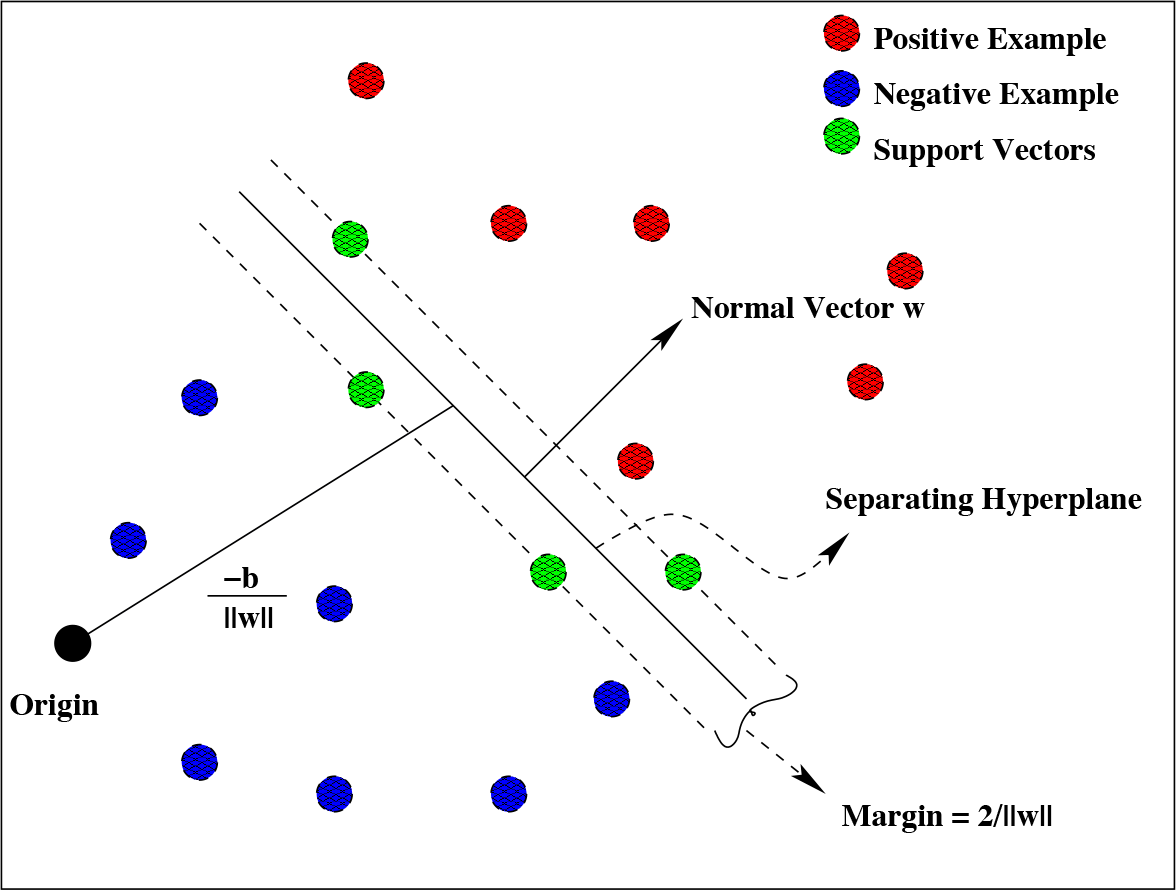
Linear hyperplane for separable data. Adapted from Burges (A Tutorial on Support Vector Machines)

#### 10.3 Lagrangian Representation: Separable Case

Clearly, the previous paragraph shows that finding the margin is a problem of optimization as the goal is to minimize ||**w**|| ^2^ subject to constraints in Eqn. 47. Employing the ideas of Chapter 3, the Lagrangian for the above problem, is:

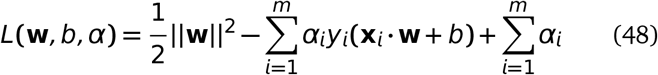

where 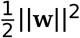 is the objective function, *α* is the Lagrangian multiplier and the Eqn. 47 is the inequality constraint. Since the minimization of the objective function is required, we employ the ideas of the derivation of KKT conditions (Eqn. 42) to Eqn. 48. In short, we would require the *L*(**w**, *b, α*) to be minimized w.r.t **w** and *b* and also require its derivative w.r.t all *α_i_*’s to vanish. Thus the KKT conditions take the form:

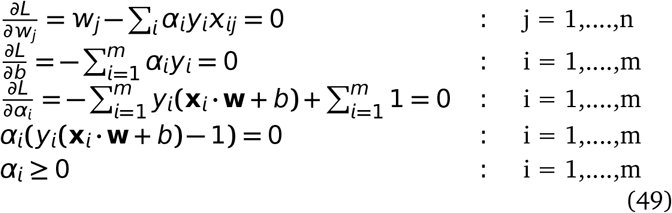

Thus solving the SVMs is equivalent to solving the KKT conditions. While **w** is determined by the training set, *b* can be found by solving the penultimate equation in Eqn. 49 for which *α_i_* ≠ 0. Also note that examples that have *α_i_* ≠ 0 form the set of support vectors.

The dual problem for the same Lagrangian is:

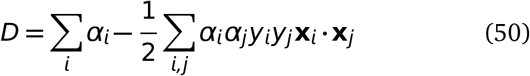

Solving for Eqn. 50 requires maximization of *D* w.r.t *α_i_*, subject to second line of Eqn. 49 and positivity of *α_i_*, with the solution given by first line of Eqn. 49.

To classify or predict the label of a new example **x**_*new*_, the SVM has to evaluate (**x**_*new*_ ·**w +** *b*) and check the sign of the evaluated value. A positive sign would lead to assignment of a **+**1 label and a negative sign to −1.

#### 10.4 Nonseparable Case

For many classification problems, the data present is nonseparable. To extend the idea to nonseparable case, some amount of cost is added, which takes care of particular cases of examples. This is achieved by introducing slack in the constraints Eqn. 45 and Eqn. 46 (Burges ^77^, Vapnik and Vapnik ^80^). The equations then becomes

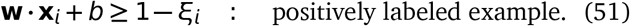

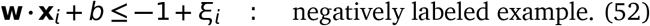

For an error to occur, the *ξ_i_* value must exceed unity. To take care of the cost of errors, a penalty is introduced which changes the objective function from ||**w**||^2^ */* 2 to ||**w**||^2^ */* 2 **+** *C*(Σ_*i*_ *ξ_i_*)^*k*^. Thus Σ*i ξ_i_* represents the upper bound on the training error. For quadratic problems, *k* can be 1 or 2.

#### 10.5 Lagrangian Representation: Nonseparable Case

Since the formulation of the Lagrangian and its dual follow the same procedure, as mentioned before, we only mention the equations. The Lagrangian for nonlinear nonseparable case is:

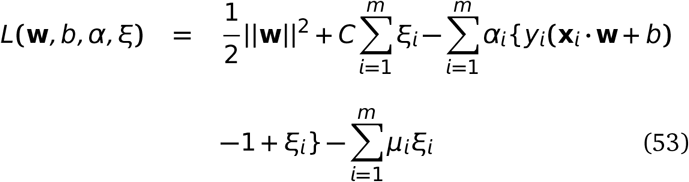

The corresponding KKT conditions are:

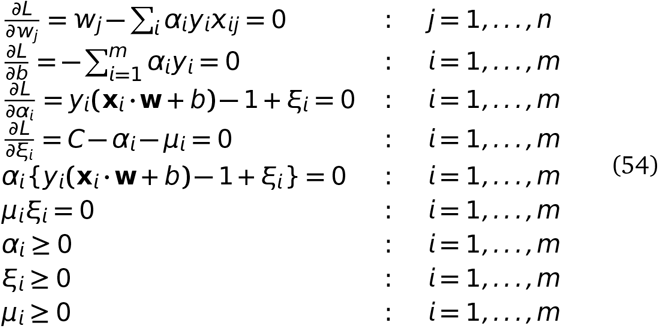

The dual formulation *k* **=** 1 for the Lagrangian just discussed is:

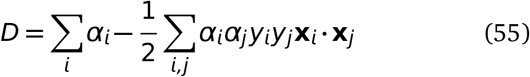

All the previous conditions remain same, except that the Lagrangian multiplier *α_i_* now has a upper bound of value *C*. The solution for the dual is given by 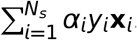. *N_s_* is the number of support vectors. Figure 12 depicts the nonseparable case.

**Fig. 12.**
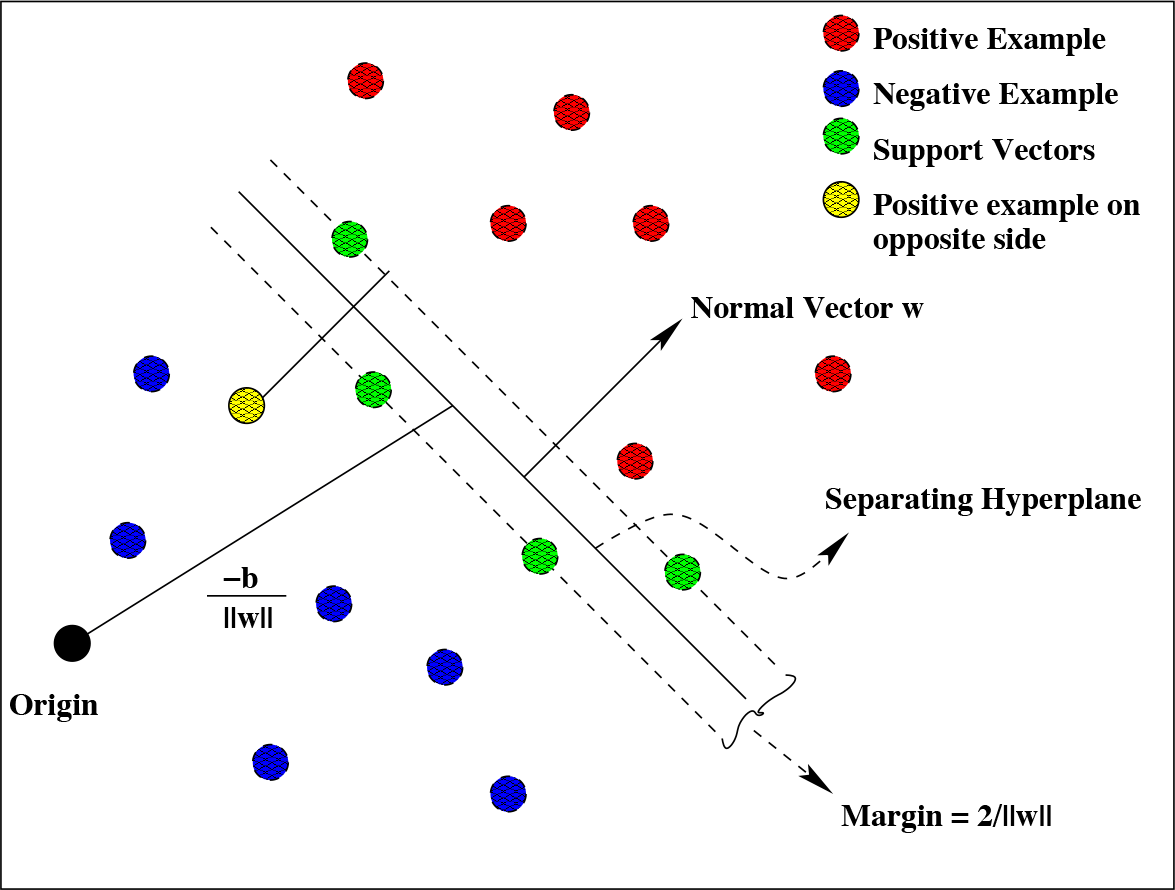
Linear hyperplane for nonseparable data. Adapted from Burges (A Tutorial on Support Vector Machines)

**Fig. 13.**
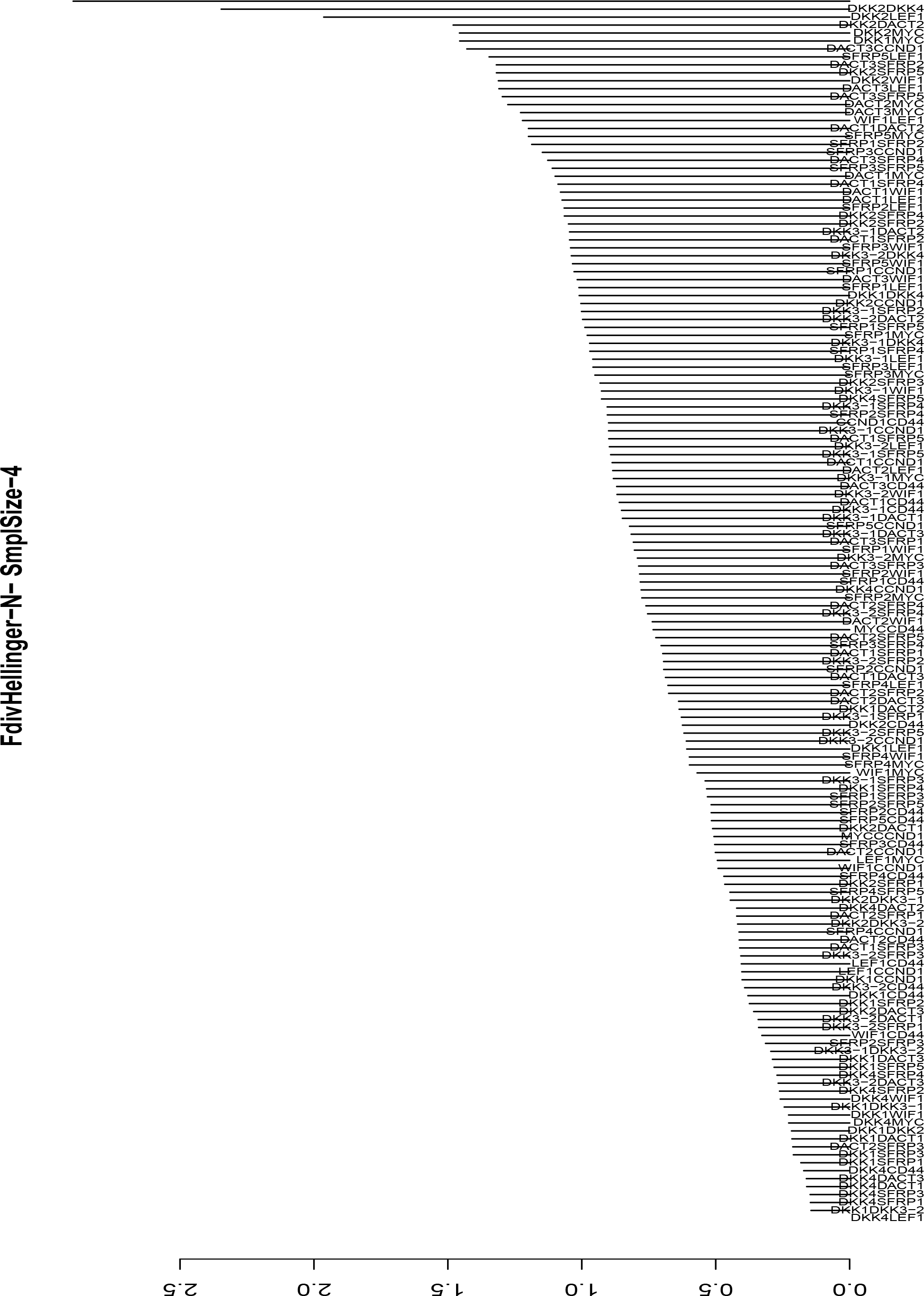
FdivHellinger; Training sample size - 4; Test sample size - 16; Case – Normal

**Fig. 14.**
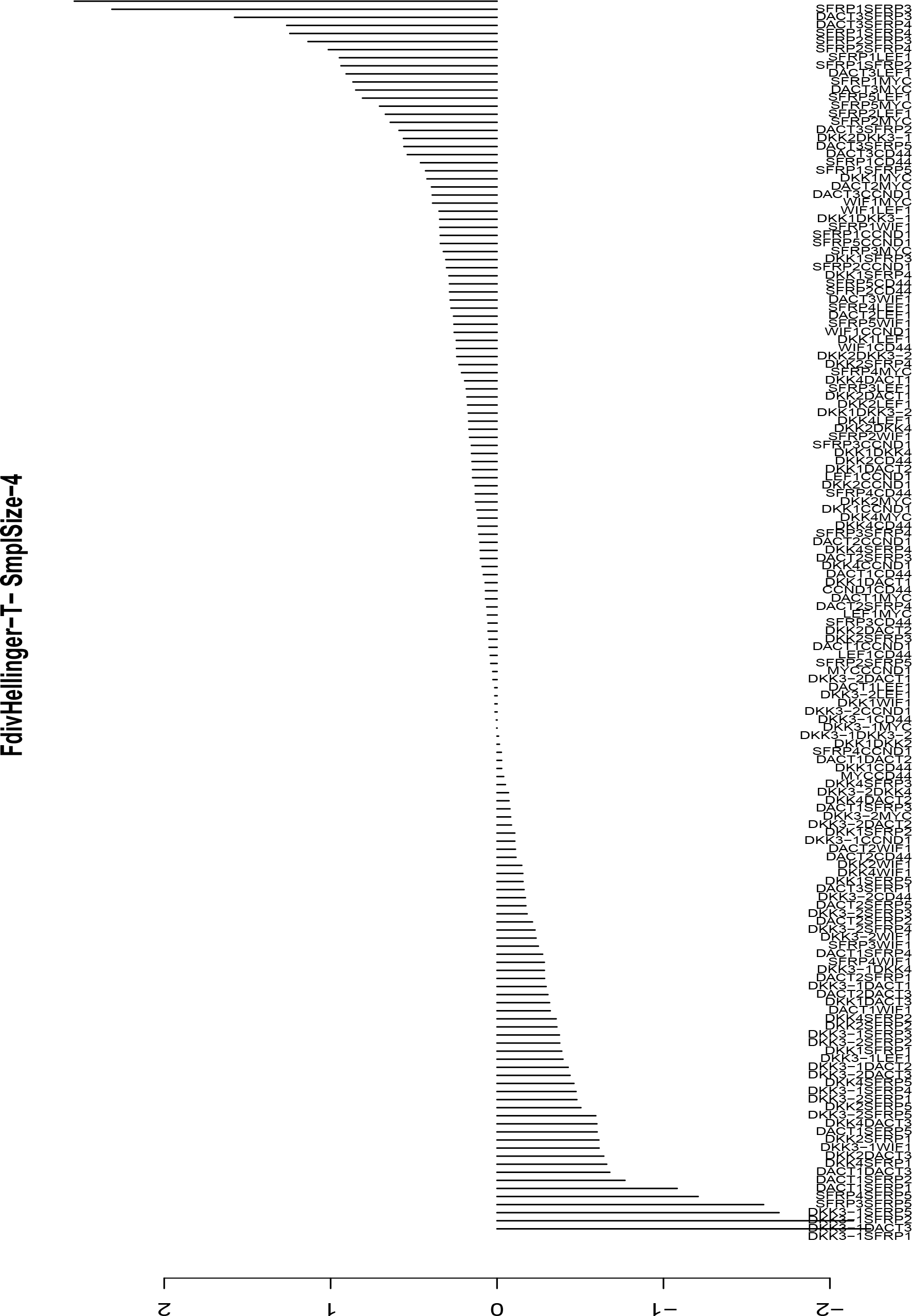
FdivHellinger; Training sample size - 4; Test sample size - 16; Case – Tumor

**Fig. 15.**
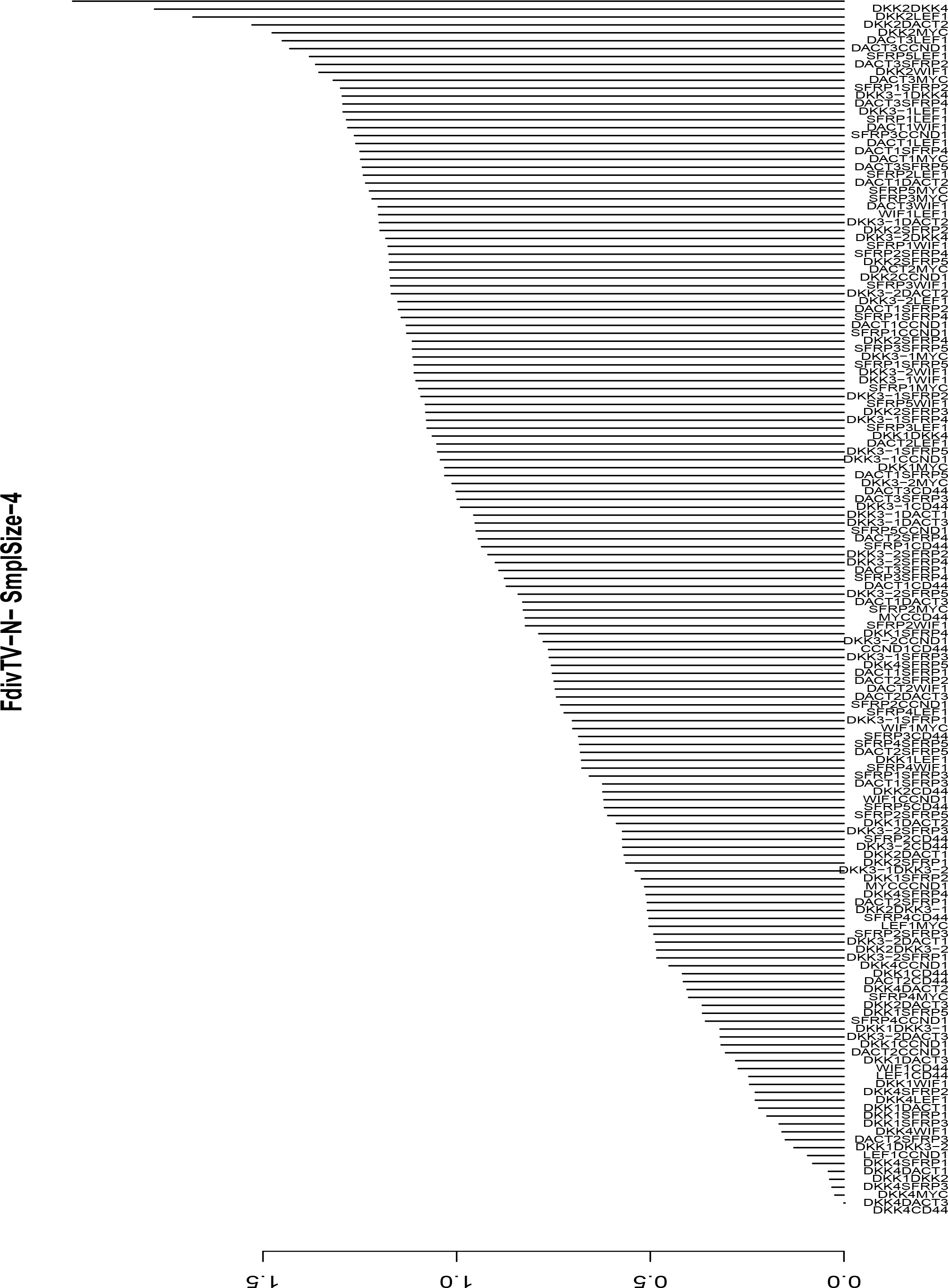
FdivTV; Training sample size - 4; Test sample size - 16; Case – Normal

**Fig. 16.**
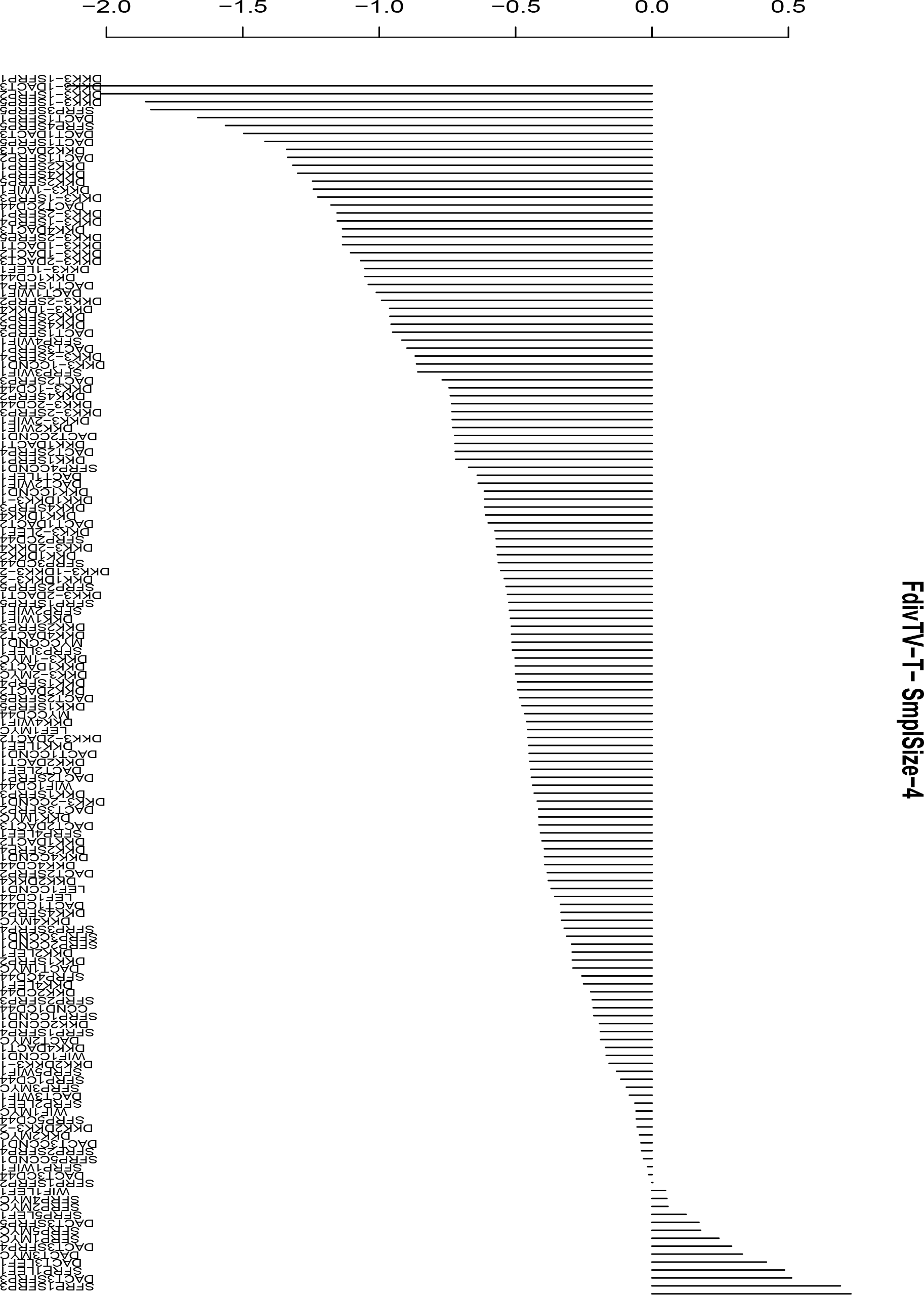
FdivTV; Training sample size - 4; Test sample size - 16; Case – Tumor

**Fig. 17.**
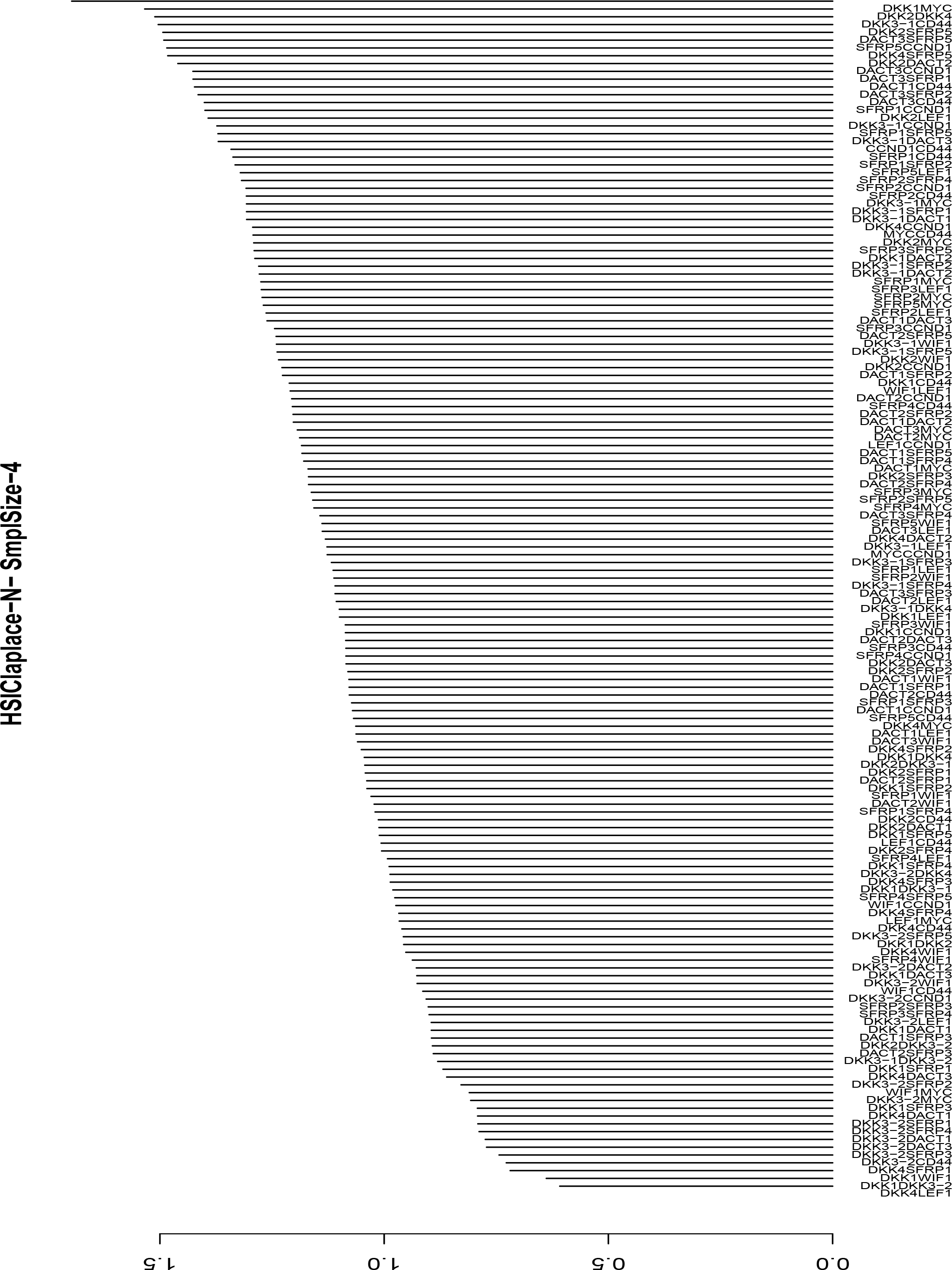
HSIClaplace; Training sample size - 4; Test sample size - 16; Case – Normal

**Fig. 18.**
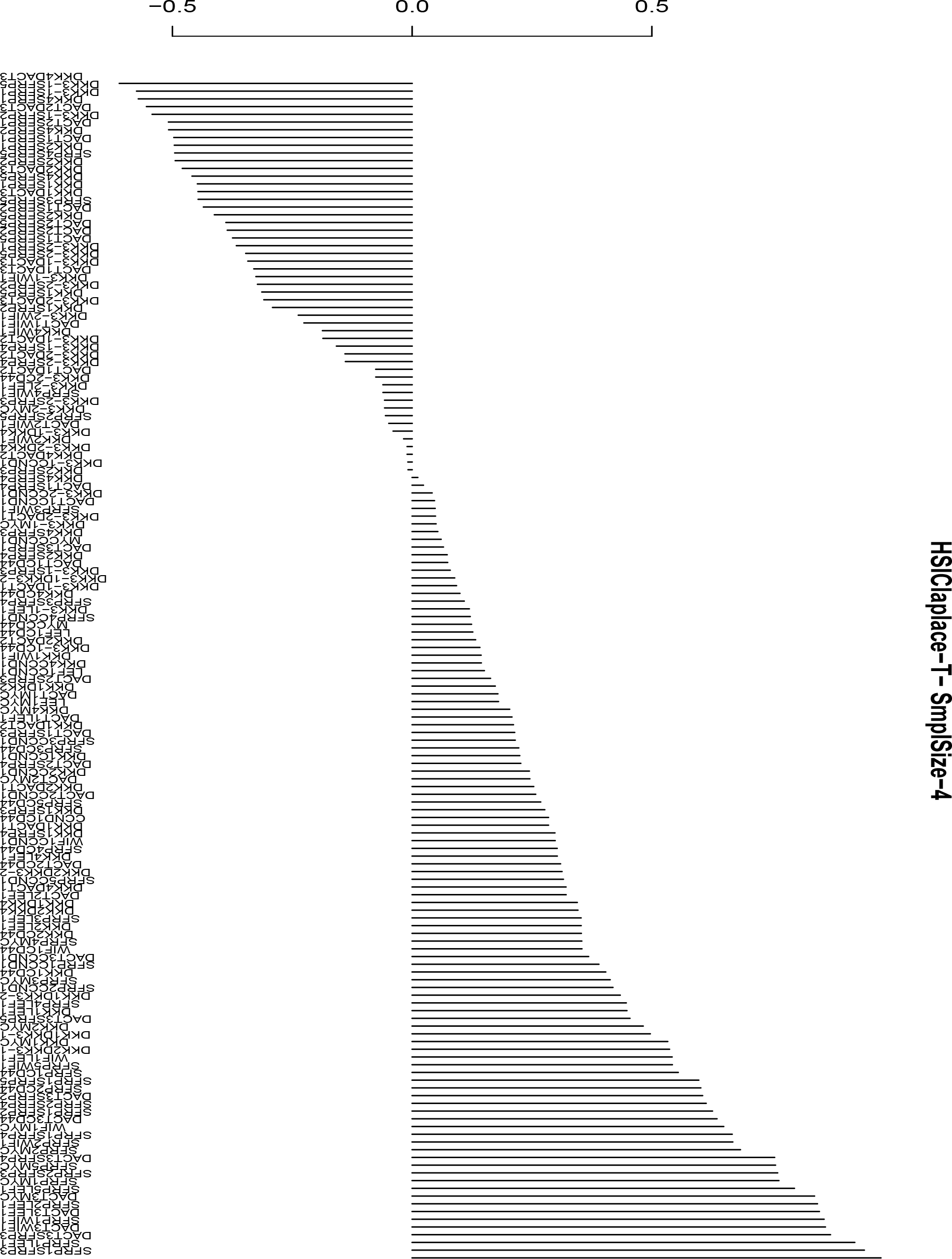
HSIClaplace; Training sample size - 4; Test sample size - 16; Case – Tumor

**Fig. 19.**
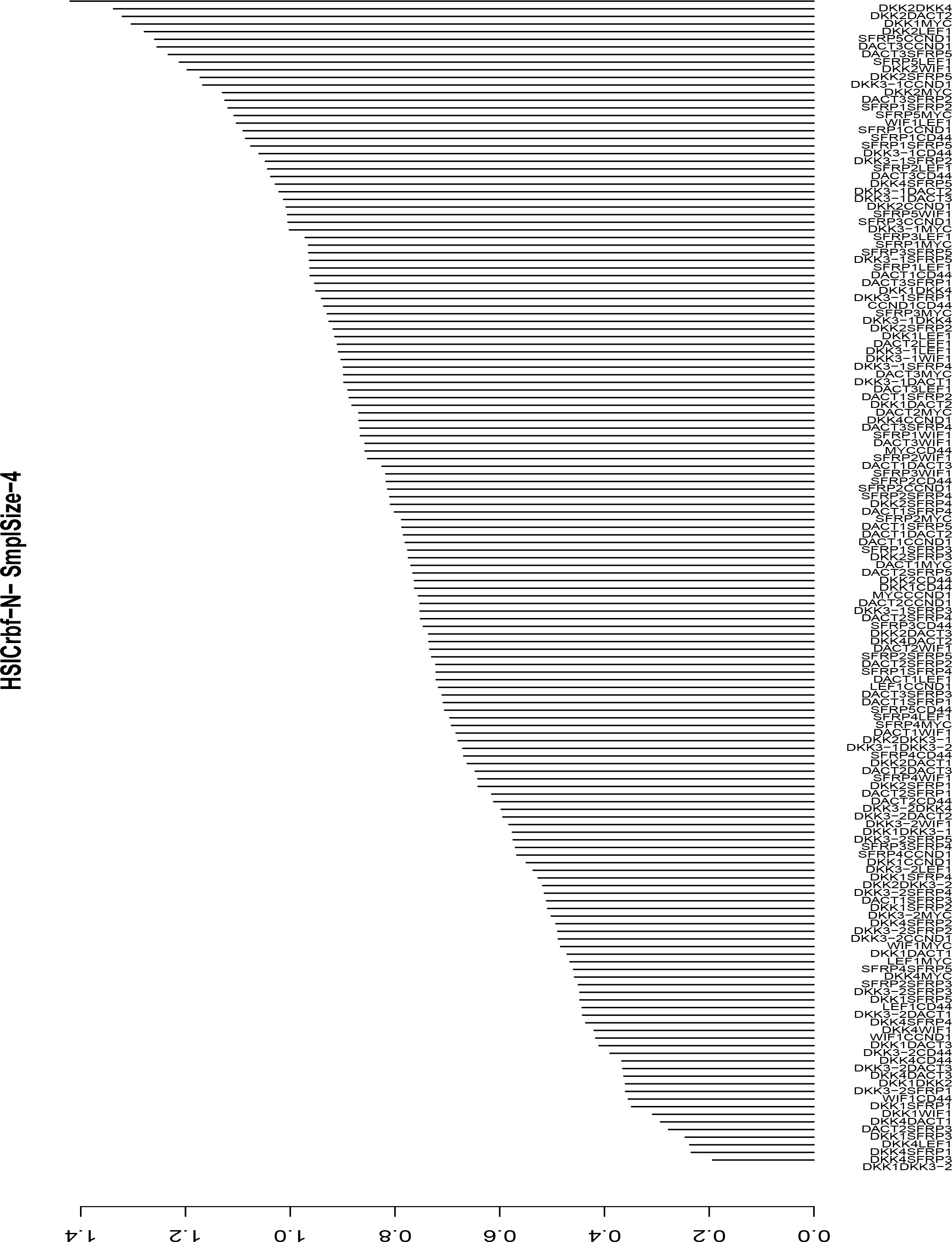
HSICrbf; Training sample size - 4; Test sample size - 16; Case – Normal

**Fig. 20.**
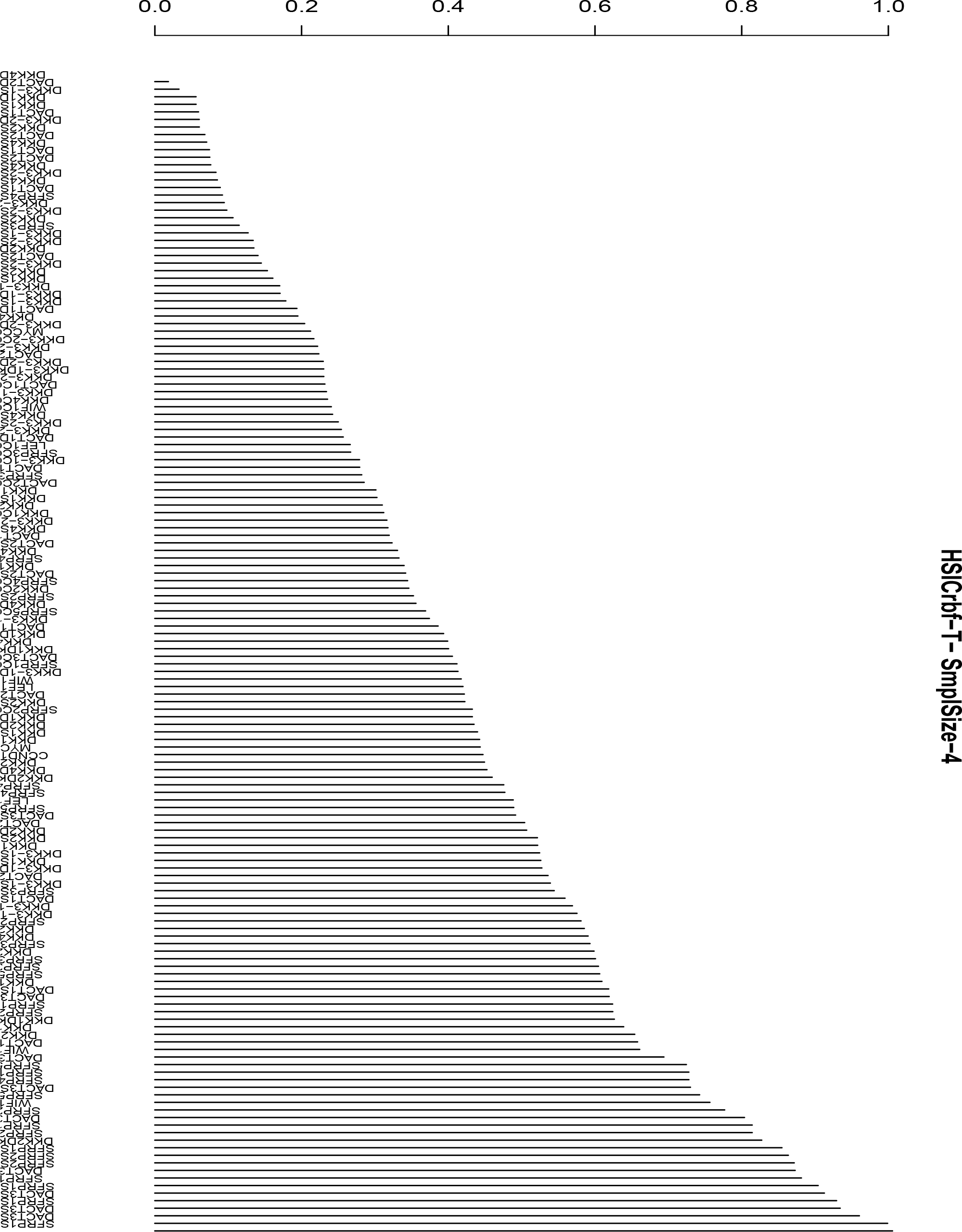
HSICrbf; Training sample size - 4; Test sample size - 16; Case - Tumor

#### 10.6 Kernels and Space Dimensionality Transformation

The above cases were for linear separating hyperplanes. In order to generalize for nonlinear cases, Boser et.al ^81^ employed the idea of Aizerman ^60^ as follows; Since the Dual in Eqn. 55 and its corresponding constraint equations employ the dot product of the examples, **x**_*i*_ · **x**_*j*_, it was proposed to map the data in a higher dimensional space using a function *ϕ* s.t. the algorithm would depend only on dot products in the higher space. Next, the existence of a function called *kernel*, dependent on **x**_*i*_ and **x**_*j*_, was assumed s.t. the value reported by the kernel was equal to the value resulting from the dot product in the higher space. A mathematical representation of the above concept is –

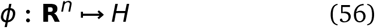

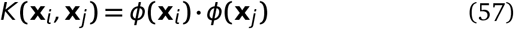

where *ℌ* is a higher dimensional space.

This technique drastically reduces the amount of work required while dealing with nonlinear separating hyperplanes, concerning search for appropriate *ϕ*. Instead, one only works with *K*(**x**_*i*,_ **x**_*j*_), in place of **x**_*i*_ · **x**_*j*_. For classification purpose, where the sign of the function (**x**_*new*_ · **w +** *b*) is evaluated, the formulation employing kernels become:

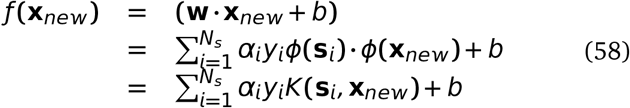

where **s**_*i*_ are the support vectors.

